# Temporal stability of functional network configurations and its relationship to cognition and psychopathology in early psychoses

**DOI:** 10.64898/2026.01.06.697919

**Authors:** Nicholas Theis, Jonathan Rubin, Ella O’Rourke, Jyotika Bahuguna, Joshua Cape, Bowei Ouyang, Satish Iyengar, Konasale M. Prasad

## Abstract

To design network-based treatments, it is critical to understand dynamic changes in brain networks over time. We comprehensively examined network structure using regional activation-only, pairwise-coactivation-only, and systems-level maximum entropy models (MEM) applied to the Human Connectome Project-Early Psychosis (HCP-EP) resting fMRI data (patients=109, controls=56). The MEM integrates regional activation levels and pairwise correlation-based functional connectivity (FC), providing the potential for better characterization of clinically meaningful brain states and how they change over time. Using the HCP/Glasser atlas to define brain regions within the default mode (DMN) and dorsal attention (DAN) networks, group differences in regional activation, graph measures of FC, and MEM features such as transition rates between minima in an energy landscape along with correlations with cognitive and psychopathological measures were examined. Psychosis was associated with reduced activation in several regions and multiple FC graph metric alterations. MEM demonstrated a wider variety of DMN and DAN activation/deactivation configurations, with more frequent switching between them, more pronounced network configuration differences and higher basin transition rates with reduced basin dwell times suggesting temporally unstable network configurations, meaning the brain cannot rely on any particular configuration for cognitive processing. Activation of only three regions correlated with working memory, but none of the FC metrics did. MEM features, in particular basin transitions and total energy, correlated negatively with working memory. Our findings suggest that MEM provides unique information, specifically attenuated inter-network connectivity with reduced stability of DMN and DAN brain states, to characterize networks as targets and their features as markers for novel treatment development.

## 1. Introduction

Network dysconnectivity has been proposed to underlie psychotic disorders[1–5]. Understanding the nature of dysconnectivity may help to elucidate pathophysiology and design novel treatments [6,7]. Existing functional magnetic resonance imaging (fMRI) studies using regional blood oxygenation level dependent (BOLD) signal properties, called first-order analyses[8,9], and functional connectivity (FC) characteristics, termed second-order analyses[6], have reported important findings [3,10–15] that may not reflect inherently dynamic functional network properties[10–13] at refined time-scales of individual repetition times (TR) or at spatial scales involving patterns across multiple regions. Cognitive processes and psychopathology involve time-variant changes in network organization, justifying the examination of dynamic network changes[6,7].

Elucidating temporally refined network dynamics is clinically significant because static FC cannot detect pathophysiology until it has progressed to the expanded time-scales at which first-and second-order analyses operate, as noted in schizophrenia [16] and synucleopathies[17]. Dynamic FC (DFC) studies using sliding windows show that schizophrenia patients spend less time in strongly and widely connected states and more time in weakly connected states with infrequent switching, associated with psychosis severity[18–20]. Alterations in connectivity may also be state-dependent rather than persistent, e.g., an association between attentional-system integration and hallucinations, or between DMN segregation and avolition, emerges only within particular dynamic configurations[16,21] suggesting that DFC analysis can disambiguate psychotic symptoms. Magnetoencephalography and fMRI data -acquired at different timescales-revealed different properties of DFC in schizophrenia[16–18]. Hence, it is critical to understand patterns of dynamic functional network changes at refined timescales.

While some DFC analyses can be superior to others[19], elucidating temporal changes relies mainly on sliding window-based DFC that necessitates window sizes of ≥10 TRs and user-defined shifting intervals that can vary across studies making it difficult to compare the findings. Such windows, shifted over multiple TRs, may not capture temporal network changes in shorter timescales nor provide a global landscape of all possible FC states and do not detect attractor states[18], i.e. connectivity patterns to which the brain frequently gravitates. The pairwise maximum entropy model (MEM) can address these shortcomings by elucidating transitions between network configuration at each TR by explicitly integrating both regional activation and pairwise correlations and by quantifying the dynamic properties of the system. MEM applied to fMRI data[20,21] captures novel pathophysiological metrics of clinical significance[22] such as “energy” values (distinct from metabolic energy) that inversely correlate with the probability of occurrence of network states defined as patterns of regional activation-deactivation. High energy network configurations occur with low frequency and hence are considered to be unstable, and time series characterized by many such states are indicative of a lack of consistency in brain activation patterns. Energy levels derived from MEM analysis also provide a basis for the derivation of “energy landscapes”[20], which represent all configurations within the state-space, identify local energy minima[23] called basins, and yield metrics like rates of basin transitions and basin dwell times. These quantifiable metrics characterize dynamic reconfigurations of systems-level networks that may underlie cognitive impairments and psychopathology.

We applied the MEM framework to the Human Connectome Project-Early Psychosis (HCP-EP) data[24] alongside regional BOLD signal comparisons and FC analysis to assess the utility of these approaches with a possibility of improving confidence in results and interpretability[25,26]. Instead of raw BOLD signals, we examined the BOLD signal frequency bands because of their better concordance with local field potentials (LFPs), especially the slow bands 4 and 5 with resting state network properties[27]. In schizophrenia, altered BOLD activity is proposed to reflect electrophysiological changes[28]. Prior studies have found altered power across specific resting-state frequency bands, reduced γ-activity, and altered neurovascular coupling in schizophrenia[29] suggesting disruptions in the timing and coordination of neuronal activity. Lower frequency bands have been associated with reduced power in schizophrenia compared to controls for DMN regions[30]. Based on the existing literature, we predicted the psychosis cohort to have altered regional BOLD signal intensity and graph metrics reflecting network dysconnectivity.

We focused on the DMN, which showed replicated evidence of abnormalities in schizophrenia[31,32]. The DMN plays a role in both resting state *and* task activity[33], and is central to many aspects of human cognition[31,32]. Another widely studied system, the dorsal attention network (DAN) may be responsible for attentional deficits considered intrinsic to schizophrenia [34–36]. While DMN may be involved in self-referential thinking, planning, retrospection, and prospection, the DAN is associated with hierarchical management of external stimuli and goal orientation[36] and the two networks are often anticorrelated[37]. Altered interaction between the DMN and DAN may explain cognitive impairments and psychopathology in schizophrenia[37].

We hypothesized that patients would exhibit more high-energy states (low probability network configurations) and more basin transitions with reduced basin dwell times based on our prior observations on adolescent-onset schizophrenia [38] compared to controls, thus highlighting the dynamic interactions within and between the DMN, DAN, and the combined DMN+DAN system. These measures reflect the temporal stability of functional network states. Few studies have examined these networks using this approach at the granularity of the HCP/Glasser atlas [39] across patients and controls.

## 2. Methods

### 2.1 MRI Preprocessing

The T_1_w structural and resting-state fMRI in the anterior-to-posterior encoding direction from the HCP-EP dataset[24] underwent the HCP minimal preprocessing [40] on the Pittsburgh Supercomputing Center’s Bridges-2 system[41]. Using FreeSurfer (v7.4.1), T_1_w images were intensity normalized, tissue labeled, and segmented [42]. The FMRIB Software Library (FSL v6.0) was used for fMRI preprocessing[43] as described previously[44]. Rigid body motion correction, head motion estimation, and gradient distortion correction (via TOPUP) were applied to each volume. Analysis of HCP-EP data was approved by University of Pittsburgh IRB (STUDY19030036).

Following the registration of T_1_w images to fMRI space and aligning the atlas, the 360-region HCP/Glasser cortical atlas[39] was applied [45] to create subject-space atlases. The DMN (34 bilateral nodes) and DAN (24 bilateral nodes) defined on the HCP/Glasser atlas are non-overlapping and were from the posterior cingulate, anterior cingulate, and lateral parietal areas[46], and the frontal, parietal, and occipital cortices [47], respectively (**Supplement 2**).

### 2.2 Regional Activation Analysis

In the frequency domain of the fMRI signal, slow oscillations (0.01-0.1Hz)[48] reflect coordinated neural activity[49] via neurovascular coupling and hemodynamic response[50], which cause local changes in magnetic susceptibility[49,51]. Five canonical frequency bands are described: slow-5 (0.01Hz-0.027Hz), slow-4 (0.027-0.073Hz), slow-3 (0.073-0.198Hz), slow-2 (0.198-0.5Hz), and slow-1 (>0.5Hz). Slow-1 overlaps with the lowest frequencies of the classic δ band and is beyond the fMRI Nyquist limit (∼0.7Hz). Slow-5 and slow-4 have been associated with LFP periodicity[52] and hemodynamic response function[53]. Often, slow-3 is partially filtered to completely exclude slow-2, which includes the respiratory frequency of ∼0.3Hz[54]. Studies that combine fMRI with electrophysiology[54] suggest high frequency LFP activity in the γ-band is resource intensive and thus more likely to increase BOLD signals [55].

For each subject and region, the spectral power of the four frequency bands was calculated, and brain-wide regional band power were compared between groups using MANCOVA controlling for age, sex, and lifetime antipsychotic exposure in chlorpromazine equivalents (LCPZE) followed by post-hoc between-subjects’ effects for group differences in each region, correcting for multiple tests (**Supplement 2**)(SPSS v32). A MANCOVA model was separately applied to slow-5 only, to compare the 58 DMN and DAN nodes by group.

Next, nodal activity was binarized to classify nodes as “on” or “off” by normalizing the BOLD signals (Section 2.4). The number of times each node switches from off-to-on or on-to-off from one TR to the next (per subject and per node), was recorded as nodal “microstate switches,” which were compared across all subjects to each frequency band’s power. We correlated switching with band power at whole-population and within-group levels. Nodes were ranked by switching frequency and top and bottom switching nodes were selected. The top-switching network was selected under the assumption that nodes exhibiting the highest switching activity are likely to be the most informative with respect to network dynamics. However, to ensure that our findings were not solely driven by this assumption or biased toward the most active nodes, we also examined the bottom-switching network, comprising the nodes with the lowest switching activity. Analyzing both extremes of the switching spectrum allowed us to assess whether the observed patterns were specific to highly dynamic nodes or reflected more general network properties. Same set of top-switching and bottom-switching nodes were used for both the groups. The group differences were assessed using Kolmogorov-Smirnov (KS) tests.

### 2.3 FC Analysis

We examined the 34-node DMN-only, the 24-node DAN-only, and the 58-node DMN+DAN built using pairwise correlation of regionally averaged fMRI timeseries [6], including edges with a correlation at α=0.05 (uncorrected) where thresholding was necessary. The Brain Connectivity Toolbox[56] was used to quantify weighted graph measures of integration, segregation, modularity, and centrality measures, namely nodal clustering coefficient (CC), modularity, and efficiency. Nodal betweenness centrality (BC) was calculated on binary graphs resulting from thresholding.

Whole-graph and nodal network measures were compared between patients and controls using MANCOVA controlling for age, sex, and LCPZE. For the whole-graph analysis, the dependent variables included average CC, global efficiency, modularity, and average BC from DMN and DAN and DMN+DAN system. Nodal-level analyses treated CC, BC, and strength as repeated measures within subjects. Post-hoc between-subjects effects analysis were corrected for multiple tests (Bonferroni). Edge-level FC was examined by comparing all unique pairwise correlations between nodes, with significance at α=0.001. The above analyses were repeated for networks constructed within individual frequency bands (**Supplement 3**).

### 2.4 MEM Analysis

Details of MEM implementation are described previously [20,22,23,38]. Briefly, the pipeline from fMRI-to-MEM identifies a parameter vector *h*, corresponding to node activation (first-order), and a parameter matrix *J,* corresponding to co-activation of node-pairs for binary network state data (second-order). To estimate these parameters, we computed a set of state vectors of dimension equal to the number of nodes in the network, one for each measured time point across all subjects, with entries 1 for active nodes and –1 for inactive. MEM parameters (*h, J*) were estimated from binary state vectors concatenated across all subjects and timepoints (165×410=67,650 observations) via a two-stage pseudo-likelihood/L-BFGS-B procedure (55 free parameters; observation-to-parameter ratio ∼1:1,230, obviating explicit regularization; full optimization details in **Supplement 4**) [57]. Parameter robustness was confirmed via permutation analyses across 500 randomly selected node subsets **(Supplement 4)**.

To compute the state vectors, each subject’s BOLD timeseries was z-scored per node and binarized at zero (active if z>0, inactive if z≤0), a threshold that maximizes marginal Shannon entropy and has been validated in prior MEM work along with other thresholding and normalization choices[22,58,59]. MEM analysis was performed separately for DMN, DAN, and DMN+DAN networks comprising brain regions (nodes) that exhibited the most frequent (top-switching) or least frequent (bottom-switching) changes in binarized state (i.e., switches between 0 and 1, corresponding to activation levels crossing the binarization threshold). The number of distinct binary states per subject for each of the DMN, DAN, and DMN+DAN networks (the DMN+DAN networks had 5 nodes from each component) was calculated and group differences in distribution were assessed using 2-sample KS tests.

After fitting MEM parameters, subject-level energy traces were computed by summing instantaneous state energies across timepoints and compared between groups using KS tests. Disconnectivity graphs were constructed from local energy minima using binary Hamming distances[23], with basins defined by monotonically decreasing energy paths from each state to its nearest local minimum. Subject-level basin transition rates and dwell times were derived from successive basin assignments and compared between groups using KS tests. Analyses used a modified version of *elapy* (https://github.com/okumakito/elapy) and custom MATLAB code **(Supplement 5)**. To evaluate generalizability of these findings across all possible 10-node MEM network configurations within the DMN, DAN, and DMN+DAN, we conducted permutation tests using 500 randomly selected 10-node sets for each system **(Figure 3; Supplenent 4**).

### 2.5 Association with cognition

Next, the association of imaging-derived variables demonstrating significant group-differences with performance were examined including attention (*d’* from auditory continuous performance task, ACPT), list sorting working memory test (LSWMT), the Flanker test for executive function, and age-adjusted dimensional change card sort (DCCS) test from the NIH Toolbox. Group differences were assessed using MANCOVA controlling for age, sex, LCPZE, and diagnostic status, followed by Games–Howell post hoc tests. Associations between cognitive performance, first-order metrics, graph measures, and MEM-derived metrics were examined using partial correlations controlling for age, sex, LCPZE, and diagnostic status, with false discovery rate (FDR) correction for multiple comparisons.

To assess whether MEM-derived metrics predict working memory beyond first-and second-order measures, we conducted hierarchical regression analyses. Because MEM takes first-and second-order statistics as inputs and derives higher-order dynamic metrics from the resulting energy landscape, it cannot be treated as a sequentially independent block alongside those measures. We, therefore, compared two complementary models predicting LSWMT performance (N=155). Model A entered variables (age, sex, LCPZE, AP/NAP) in Block 1, followed by a first-order band power composite (PC1; 79% variance) in Block 2, and a second-order FC graph metric composite (PC1; 88% variance) in Block 3. Model B entered the same variables in Block 1, followed by a MEM composite of five basin-transition and energy-trace metrics significantly associated with working memory (PC1; 66% variance) in Block 2. PCA composites were used to address collinearity among raw FC metrics (r=0.99; VIF>100). A sensitivity Model C additionally tested sequential entry of covariates, MEM, first-order, and second-order FC **(Supplement 6)**.

## 3. Results

### 3.1 Sanple Characteristics

The sample consisted of 165 subjects (patients=109, non-affective psychoses (NAP)=76%, affective psychosis (AP)=24%; controls=56) (other HCP-EP studies included 93-124 patients[60–62]). Patients were younger than controls with no sex distribution difference **(Supplement 7)**. Mean framewise displacement of rigid body head motion for resting state did not show group-wise significant differences (controls=1.35±0.93; patients=1.25±1.04; t=0.61, p=0.55).

### 3.2 Regional Activations

At the whole-brain level, MANCOVA models after controlling for age, sex, and LCPZE were not significant for any band. Post-hoc tests of between-subjects’ effects with Bonferroni correction showed differences in ≈10% of regions in BOLD signals and were unique to each band; ≈50% showed decreased BOLD responses while the rest showed elevated BOLD responses. Within DMN and DAN, 3 of the 58 regions showed altered BOLD frequency power (MANCOVA; Wilk’s λ=0.52, F=1.63, p=0.016; post-hoc between-subjects’ effects, p_Bonferroni_ all<0.05) (**Supplement 2**). The power in every band negatively correlated with microstate switching **(Fig 1A)**.

**Figure 1:**
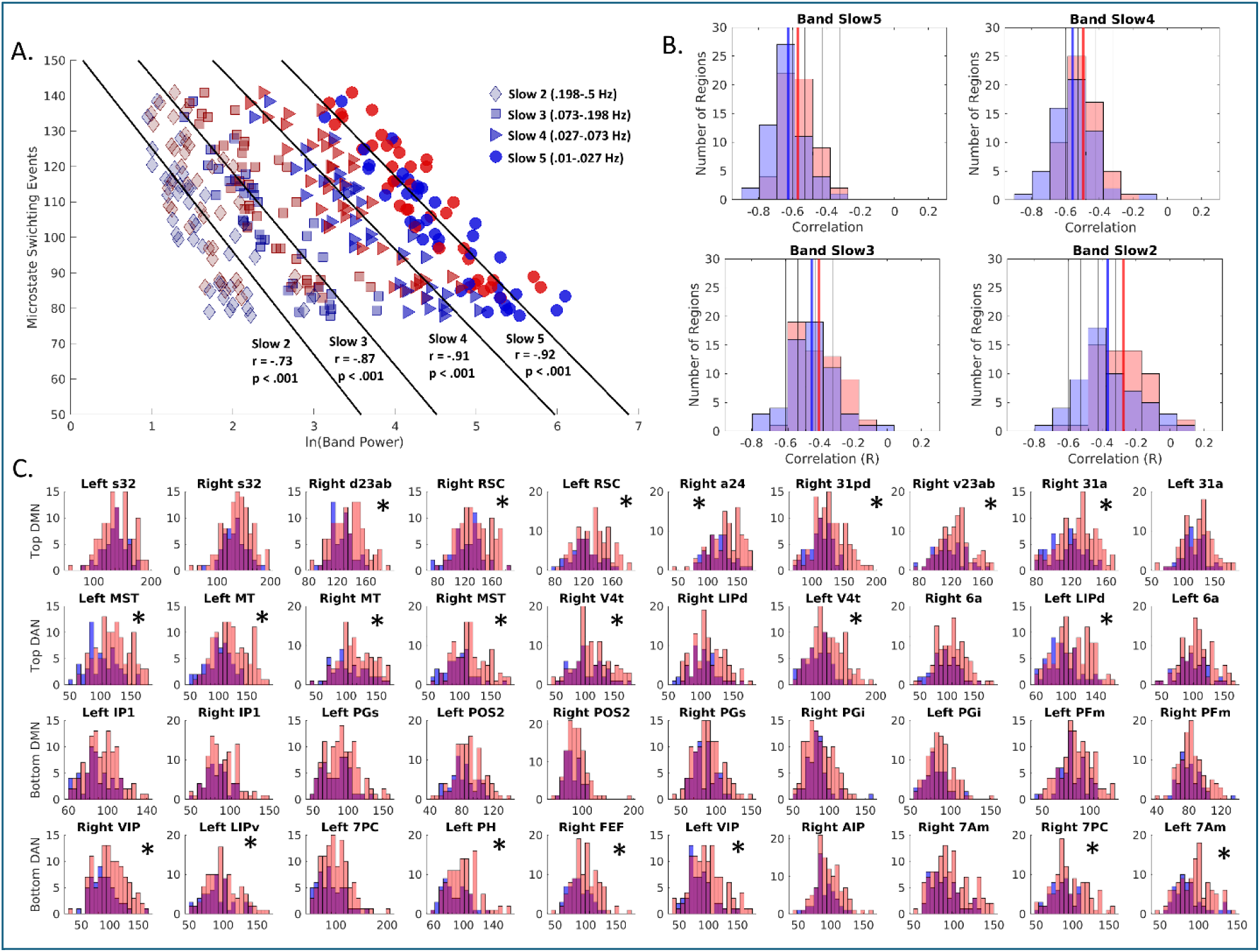
First Order Results showing group-level average band power versus the number of region-wise microstate switching. **A)** Correlations between power in frequency bands and counts of microstate switching events, where each scatter point represents the group-level average (controls in blue, patients in red) for a region. **B)** Distributions across subjects of correlations between switching and power, for each band, by group. Black vertical lines show band averages from all bands (including sibling subplots), whereas blue and red vertical lines show control and patient (respectively) group level average correlations (of regions) for that subplot. Histograms show nodal distributions of correlations calculated across subjects within a group. **C)** Microstate switching by group: asterisks indicate significant differences in patient-control distributions by KS-test after FDR correction at level alpha = 0.05. For both DMN and DAN, the top (most microstate switching events) and bottom (least microstate switching events) 10 regions are shown. Histograms represent all individuals, where y-axis is number of participants and x-axis is number of microstate switching events in the given region over the course of the resting state fMRI acquisition.

### 3.3 FC Results

We observed a non-significant group main effect on the combined global network measures controlling for age, sex, and LCPZE (MANCOVA, Wilks’ λ=0.99, F(4,157)=2.32, p=0.059). Post-hoc tests showed significant group differences in all global graph metrics (p_Bonferroni_<0.02) (**Supplement 3** for graph metrics of DMN, DAN, and DMN+DAN) but nodal graph metrics did not show case-control differences (MANCOVA, Wilks’ λ=0.99, F(2,160)=0.05, p=0.95); post-hoc Bonferroni-corrected tests showed higher BC (p=0.011), lower CC (p=0.015), and strength (p=0.0079) in patients compared to controls.

### 3.4 MEM Results

The fitted MEM energy functions for all systems matched the empirically observed state probabilities (**Supplement 4**) and predicted underlying probabilities of occurrence of each possible state in all systems. In all six systems, patients showed significantly more unique states (on/off node patterns at each TR) per subject than controls (**Figure 2A**).

**Figure 2:**
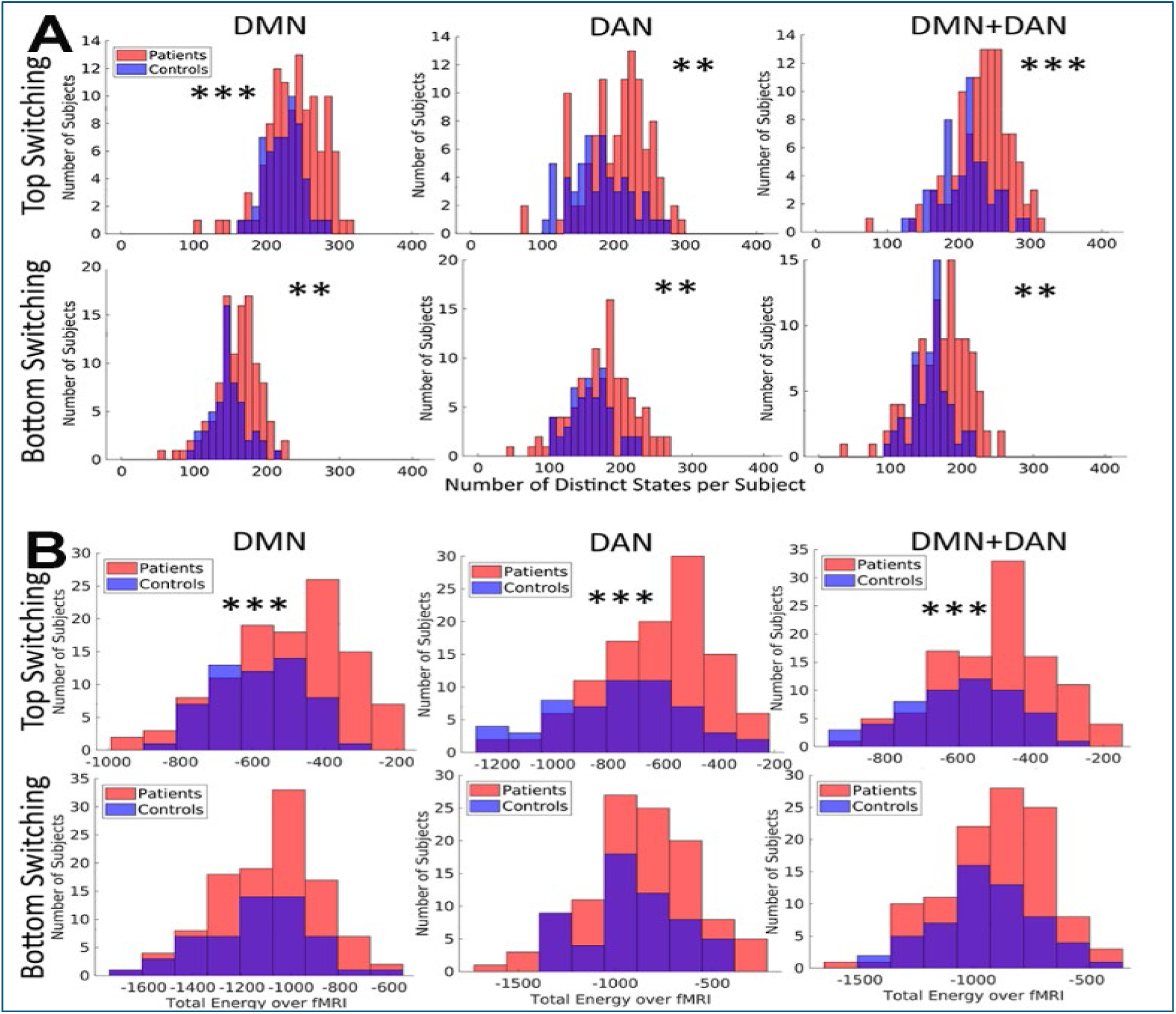
A. The number of distinct states per subject is shown in two histograms (one per group) for the six systems (top-switching. -- DMN: KS=0.32, p<0.001; DAN: KS=0.30, p=0.002; DMN+DAN (KS=0.37, p<0.001); bottom-switching -- DMN: KS=0.31, p=0.001; DAN: KS=0.30, p=0.003, DMN+DAN: KS=0.30, p=0.001). Asterisks indicate significance level of difference of distribution by two-sample KS-test, where: **p <0.01; ***p <0.001.

**Figure 2B.** Histograms of energy traces for individuals in each group and each network, defined by the total energy values across all observed states per subject. The figure shows that the patients have higher traces than controls in the DMN-top (KS=0.29, p=0.003) DAN-top (KS=0.34, p<0.001), and DMN+DAN-top (KS=0.33, p<0.001) systems. Two-sample KS-test: **p<0.01; ***p<0.001.

Patients had more distinct states in 99% of permutations, higher energy trace in 96%, and more basin transitions in 96% for the DMN. For the DAN, 96% showed more distinct states, 95% had higher energy trace, and 77% of permutations had significantly more basin transitions in patients. For the combined DMN+DAN systems, 96% showed more distinct states for patients, 95% higher energy trace, and 96% had more basin transitions. This corresponds to the results of the model with our initial selection of nodes **(Figure 3, Supplement 4**).

**Figure 3:**
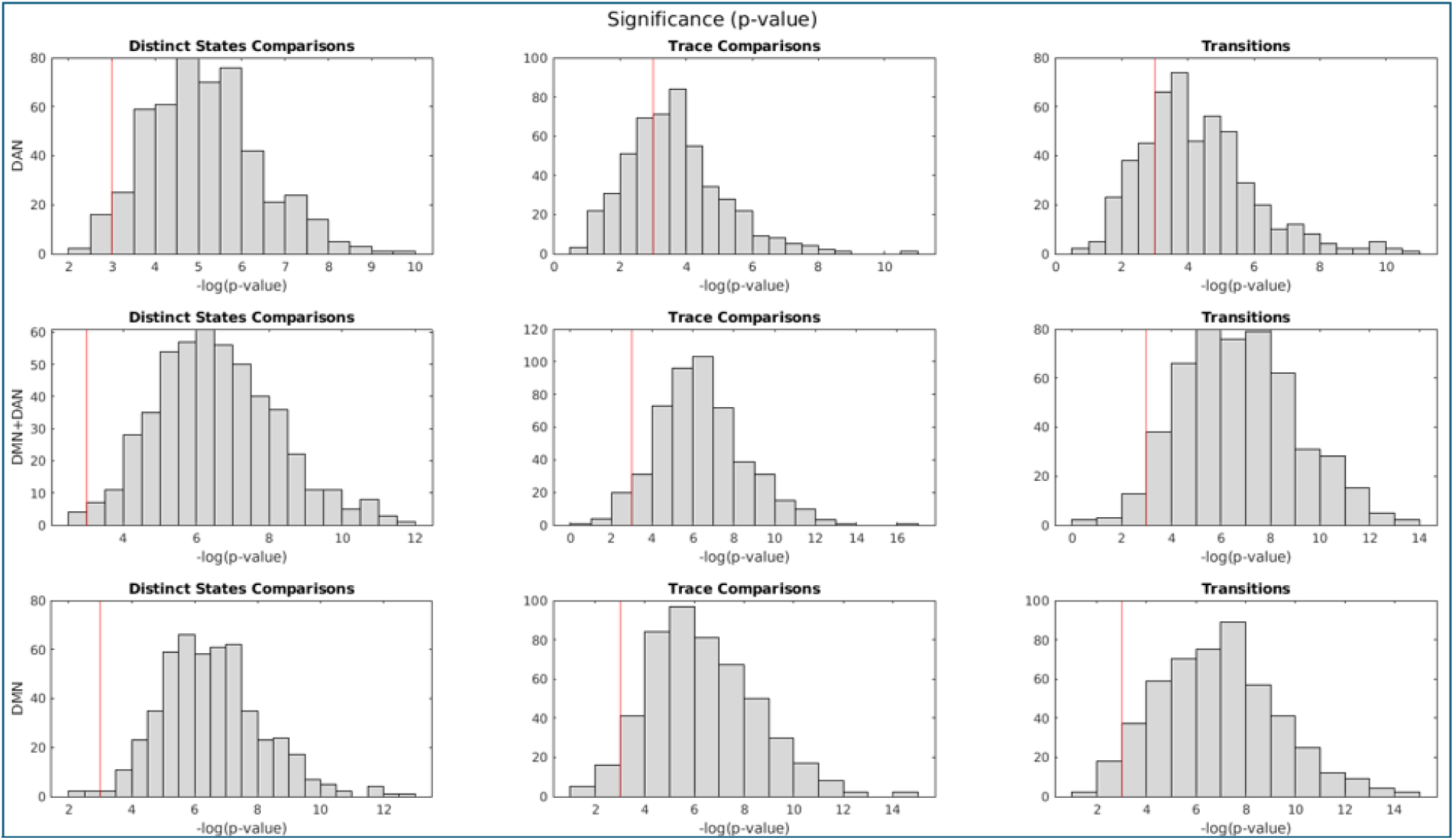
Permutation test results: Above histograms show, across the nine subplots, the rows representing the systems, namely, from top to bottom: DAN, DMN+DAN, and DMN. Columns represent summary measures, which are, from left to right: distinct states, trace, and transitions. Histograms represent 500 permutations, showing the negative log likelihood of the p-value, with the red vertical line indicating the corresponding value of alpha=0.05, and more significant values lie to the right of the red line (See Supplemental Information 5 for the plots for KS statistic values).

Between NAP and AP, no system showed significant differences in the numbers of distinct states. But NAP showed significantly more distinct states in every system, while AP showed more distinct states than controls in DMN-bottom, DAN-top, and DMN+DAN-top than controls (**Supplement 8**).

Significant patient–control differences in total energy for top but not bottom systems was observed (**Figure 2B**) mirroring first-order results, where group differences were present in top-switching but not in bottom-switching DMN nodes. Energy trace distribution was not significantly different between NAP and AP in any of the six systems, though NAP had higher energy traces in DAN-top and DMN+DAN-top compared to controls, while AP showed higher energy traces in DMN-DAN-top systems than controls (**Supplement 8**).

MEMs fit to each network of combined patient and control data yielded a shared energy landscape on which all subjects’ state trajectories were compared (**Figure 4**). The bottom-switching DMN exhibited a complex landscape, whereas the top-switching DMN had only two basins; the DAN showed an opposite pattern. Across all six systems, the all-on and all-off basins were the deepest (most stable or most visited). In the combined DMN+DAN networks, additional basins corresponding to DMN-only and DAN-only activation indicate dynamic alteration between co-activation, co-deactivation, and mutually exclusive states possibly due to anti-correlation between DMN and DAN (**Figure 2**).

**Figure 4:**
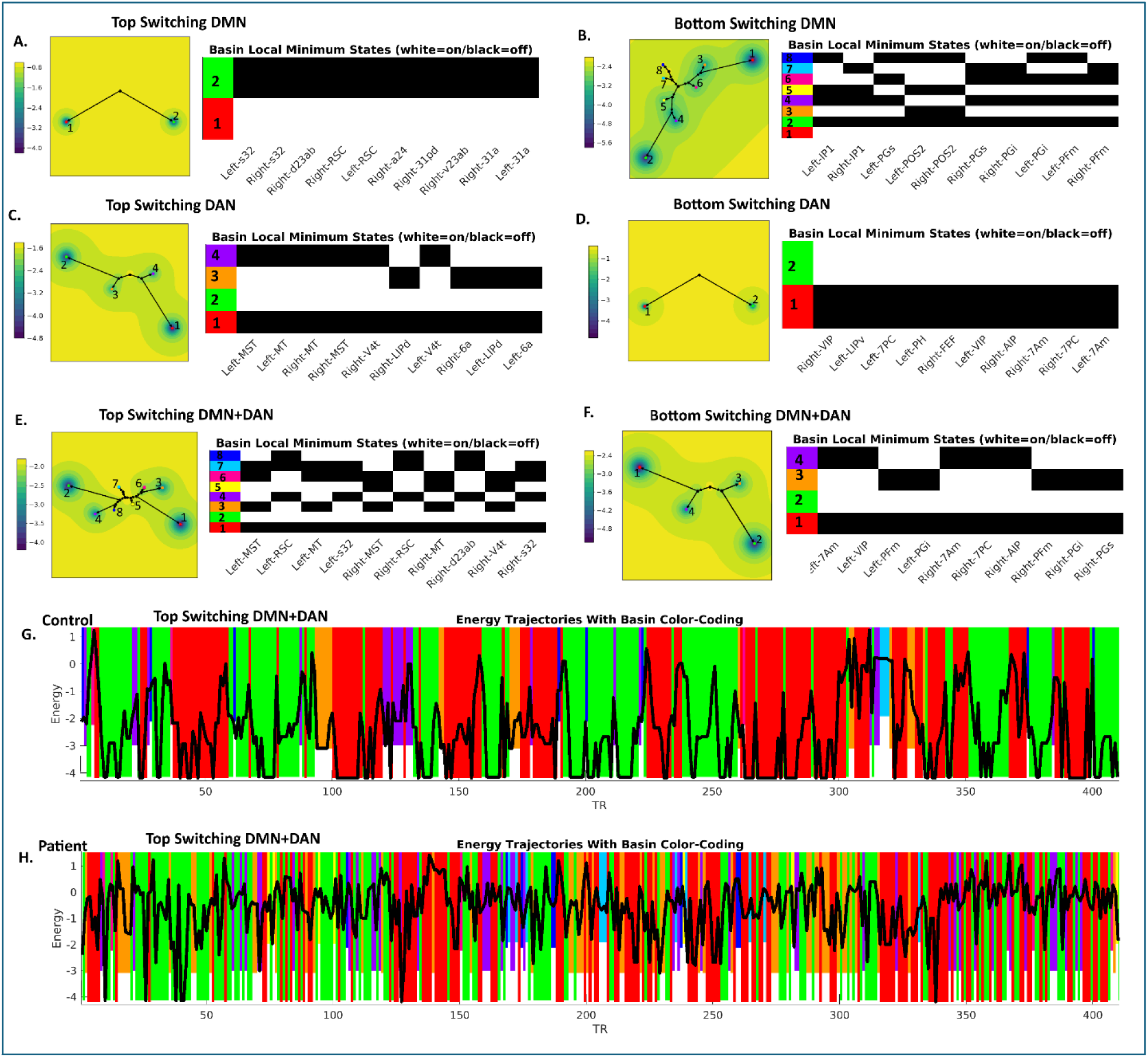
Energy ninina and basin switching within energy landscapes. **A.** Energy landscapes including minima and associated basins (left) and activation state configurations for minima (right) for top switching DMN. Analogous plots are shown for: **B.** Bottom switching DMN, **C.** Top switching DAN, **D.** Bottom switching DAN, **E.** Top switching DMN+DAN, **F.** Bottom switching DMN+DAN. **G**. The color-coded (by basin) energy trajectory for the 410 fMRI timepoints in an example control subject. **H.** The color-coded (by basin) energy trajectory for the 410 fMRI timepoints in an example patient. Both **G** and **H** use the basin color coding given in **E** and present results from an example participant’s top switching DMN+DAN. Thick black line represents energy value; each colored rectangle represents occupation of a corresponding basin; downwards extent of rectangles represent basin depth. Note that the specific state associated with each color does not agree across systems, but colors are assigned in the same order for all systems (e.g., red always identifies the system’s lowest energy basin, followed by green, and so on).

We observed significantly more state transitions in patients than controls for top-switching DMN, DAN, DMN+DAN, and bottom switching DMN+DAN (KS=0.33, p<0.001), but the fractions of transitions between specific basins were the same (**Figure 5A**) (**Supplement 6**), indicating that patients had *more* basin transitions but *not different* temporal transition patterns. In the DMN+DAN networks, the lowest-energy basins corresponded to DMN-only or DAN-only activation (basins 3 and 4 in **Figure 4E,F**).

**Figure 5:**
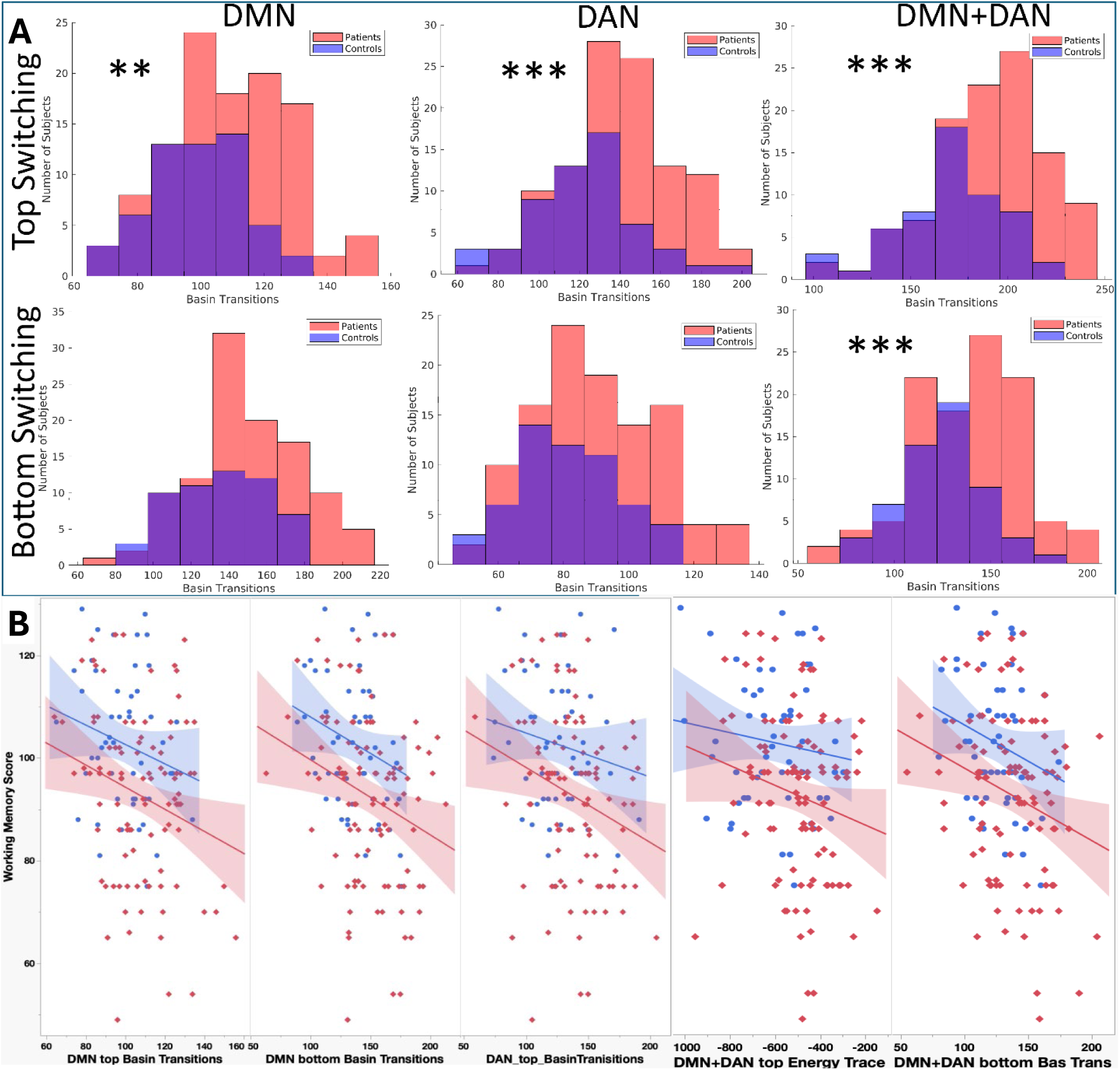
Numbers of basin transitions per subject by systen and group. Top row indicates top microstate switching networks, bottom row indicates bottom microstate switching networks. From left to right: DMN (KS=0.30, p=0.002), DAN (KS=0.32, p<0.001), DMN+DAN (KS=0.36, p<0.001). Asterisks indicate significance level of difference of distribution by two-sample KS-test: **p <0.01; ***p <0.001. **B. Scatterplot of correlations of working nenory scores with basin transitions and energy trace.** Red diamonds and fit line represent scores for early psychosis patients and blue dots with blue fit line represent healthy controls. Partial correlation controlling for age, sex, LCPZE, and diagnostic status was conducted followed by correlations for each group. All correlations for the entire sample of patients and controls were significant after correcting for multiple tests using Bonferroni approach. Groupwise correlations showed that negative correlations retained significance after multiple test corrections with the MEM metrics, but healthy controls showed significant correlations with DMN top and bottom basin transitions and DMN+DAN bottom basin transitions but not with DAN top basin transitions and DMN+DAN top energy trace.

Direct transitions from DMN-exclusive states to DAN-exclusive basins (or vice versa) were minimal (2-4%) compared to ≈40% staying in an exclusive basin, or transition from the all-nodes-on to all-nodes-off basin (∼15%), suggesting that DMN and DAN nodes switch largely independently during rest. Moreover, in the top-switching DMN+DAN system, basin 1 (all-off) and basin 3 (DMN-only) had shorter dwell times in patients, while in the bottom-switching DMN+DAN, patients had shorter dwell times in the all-on and DAN-only basins (2 and 3, respectively). Finally, NAP had more basin transitions in every system but not AP compared to controls(**Supplement G**).

### 3.5 Associations with cognitive functions and psychopathology severity

We observed a significant group main effect across cognitive measures (MANCOVA controlling for diagnostic status, LCPZE, age, and sex: Wilk’s λ=0.66, F(14,280)=4.60, p<0.001). Post-hoc Games-Howell tests (since equal variance assumption was violated) showed significant group differences for *d’*, age-adjusted DCCS, age-adjusted Flanker score, and total LSWS score (all p<0.001). NAP showed poorer performance than controls but not AP on all significant measures; only DCCS differed between AP and NAP groups.

Partial correlations controlling for above covariates showed that band power 5 in right IP1, right PFm, right PGs, right 31pd, bilateral LIPd, bilateral PH, were positively correlated with working memory (r=0.29-0.32; all p_FDR_<0.00017) and no associations with psychopathology severity. Graph metrics were not associated with both clinical measures. Several MEM-derived measures were negatively correlated with working memory after FDR correction, including basin transitions in DMN top and bottom networks (r=−0.26 and −0.27) and DAN top networks (r=−0.21), total energy in DMN+DAN top networks, and basin transitions in DMN+DAN bottom networks (r=−0.22 each, all p<0.004) **(Figure 5B) (Supplement 6)**.

To quantify the incremental predictive value of each method, hierarchical regressions with LSWMT as the outcome (N=155) showed that first-order band power explained significant variance beyond covariates (age, sex, NAP/AP, LCPZE) (ΔR²=0.075, p<0.001), while graph metrics did not (ΔR²=0.002, p=0.576; Model A R²=0.290). In the second model, MEM composite explained significant incremental variance (ΔR²=0.031, p=0.014, β=−0.183; R²=0.245), accounting for 84.4% of the variance explained by both conventional domains combined despite using one fewer predictor. A third sensitivity model confirmed that first-order band power contributed complementary variance beyond MEM (ΔR²=0.047, p=0.002), whereas second-order FC added nothing (p=0.981), indicating MEM subsumes second-order predictive content while partially overlapping with first-order measures **(Supplement 6)**.

## 4. Discussion

We investigated regional brain activation patterns using first-order, second-order, and combined MEM analyses on fMRI data from early psychosis patients and controls that provided unique insights with each method. Patients showed altered band power across multiple regions[30], replicating prior amplitude of low frequency fluctuations (ALFF) findings [8,63,64] and extending them by linking reduced band power to increased microstate switching. Association of each frequency band to different sets of regions is intriguing and may be due to local circuit architecture[65], regions operating at different intrinsic timescales[66], different structural connectivity [67], or metabolic constraints[36]. However, lack of correlation of regional BOLD signals in any frequency with clinical variables suggests the limitation of using first-order metrics alone.

Our study extends reduced brain-wide FC in psychosis [32] to more anti-correlated DMN-to-DAN connections, weaker connectivity within DMN and DAN, lower CC and efficiency, and higher modularity and BC indicating greater segregation with weaker local cohesion. Graph theoretic metrics of FC did not correlate with clinical variables either[31,68] suggesting the need for integrative methods such as MEM.

MEM formalizes system interactions consistent with neuroscience principles. [22,38,59,69–76] Though formulated in statistical physics, MEM has since been adapted across neuroscience—from retinal and hippocampal circuits to fMRI [73–79]. Pairwise MEM is a generalization of the classic Ising model [77] into higher dimensional space [78], modeling the likelihood of discrete states based on BOLD signal properties. The MEM incorporates the interdependence of pairwise correlations, unlike in FC analysis where correlations are assumed to be independent. Further, MEM can predict higher order correlations nearly perfectly (r>0.98 at every order of correlation up to the 9^th^ order) and also provide added value in terms of qualitatively different network features that can be quantified (**Supplement 10**). Energy landscapes, which represents global statistical structure of activation patterns over the entire length of fMRI data, provide novel quantifiable measures, namely energy minima (basins), dwell times, and basin transitions. Whereas traditional FC and first-order analyses describe correlation strength or amplitude of BOLD signals, MEM quantifies the extent to which systems-level brain states are stable or transient, where stable networks occur more often. In this study, patients exhibited more higher-energy states (occurrence of low probability systems-level temporal activation/deactivation pattern in the networks), higher basin transition rates, and shorter basin dwell times suggesting increased dynamical volatility of brain networks that may result from, but goes beyond, observations of altered connectivity strength alone. These MEM-derived features were not correlated with lifetime anti-psychotic exposure and were driven mainly by the NAP suggesting that some MEM features may be relatively specific to NAP. Thus, network instability, not simply dysconnectivity, may be more relevant to the neural dynamics of psychosis.

Further, MEM may account for variance beyond first-order and second-order terms in their correlation with working memory performance. This integrative property is also reflected in working memory prediction: the MEM composite accounted for 84.4% of the variance explained by the full conventional model while using one fewer predictor and without redundancy with second-order metrics (Section 3.5). This is because MEM integrates first-and second-order information within single generative framework characterizing the temporal stability and switching dynamics of network configurations and have no equivalent in first-order or second-order analyses further justifying the usefulness of MEM. As an example, MEM infers the underlying “social rules” that determine who attends a party, rather than simply observing that certain individuals attend the party more than others (first order statistics) or two people often attend the party together (second-order statistics). An example of a “social rule” could be people tend to join events attended by close friends while avoiding events attended by rivals. These inferred rules can then predict the likelihood of social gatherings that were never directly observed. Due to these unique advantages and added value, application of MEM to psychiatric research on autism to Alzheimer’s disease is gaining traction[22]. Our systematic review suggested that psychiatric disorders may exist on a continuum of network instability-to-rigidity[22]. Among three published studies on schizophrenia, one showed greater disruption to brain dynamics in schizophrenia along with increased occurrence of DMN suppression with visual activation compared to depression and controls[23]. Our studies on adolescent-onset schizophrenia task fMRI data showed greater occurrence of high energy states that correlated with executive functions [38] and reversal of dynamical complexity when patients shifted from rest to executive function task in the DMN[79]. Clinical high-risk individuals showed increased duration and occurrence of minor states with greater transitions[80]. Thus, examination of more refined temporal network dynamics can be potentially useful to better understand pathophysiology and design better treatments.

The DMN [81] and DAN [82,83] have been well-established in humans and in animal models [84], and are evolutionarily conserved[85]. Their interaction may support top-down attention and balancing of internally and externally focused thought. During rest, DAN activity may help regulate which internal representations are prioritized, while the DMN may focus on which stimuli merit attention [86]. Disruption of such interaction could contribute to cognitive impairments[87], delusional thinking [88] and thought disorganization [89]. Our results show that changes in DMN and DAN activation are mostly independent at rest, but it is likely that rest-to-task switch could show distinctly different dynamics in these networks[79]. MEM metrics correlated with cognitive impairments but not with psychopathology. This may be due to milder psychopathology (PANSS total=49.59±11.18) and medication treatment. Since cognitive impairments are proposed to as core deficits of schizophrenia[90,91] and treatment of cognitive impairments can provide better long-term improvement[92,93], correlation of MEM metrics with cognitive impairment may provide leads for a better understanding of the disorder for eventual treatment development. Approaches including MEM alongside conventional regional and network analyses can identify spatially and temporally resolved alterations in brain state dynamics linked to cognitive impairment. Future treatment development studies can potentially use these features as targets to restore them to the patterns observed in controls.

Limitations of our study are that the HCP-EP is diagnostically heterogeneous including both affective and non-affective psychosis with varying medication doses. Our cross-sectional design may not clarify ongoing pathophysiology. MEM is limited by the number of nodes that can be included because the number of possible states increase exponentially with the number of nodes, requiring very large data samples; however, we have attempted to address this by using permutation tests. Binarization of BOLD signals may result in loss of information, but prior studies show that the limitation due to binarization is minimal[20,73]. And lastly, MEM estimates the probability distribution over system states over time without explicitly modeling their temporal evolution. Consequently, while the MEM captures the relative likelihood and interaction structure of different states, it does not preserve information about the temporal ordering or transition dynamics between them. Strengths of our study include the use of state-of-the-art method to examine more refined temporal network changes at systems-level which cannot be examined using traditional DFC analysis, greater attention to motion correction and other signal artifacts, and inclusion of first-and second-order models to gain a comprehensive understanding. Our results demonstrate the distinct advantages of MEM analyses to reveal unique dynamic network features in patients, which represents a promising focal point for future studies. Future studies should extend this framework to task paradigms, longitudinal studies of network dynamics, other stages of psychosis, and treatment-related changes in neural dynamics[22].

## Acknowledgenents

We thank the NIMH NDA for providing the HCP-EP data for this research study.

## Author contributions

Nicholas Theis implemented the MEM model, analyzed the first-order and second-order data, helped model interpretation, drafted and edited the manuscript, created the figures and tables.

Jonathan Rubin helped interpret the MEM model, helped improve the model specifications and implementations, and edited the manuscript.

Ella O’Rourke implemented the statistical analyses for the first-order and second-order data, helped tabulate the results, created tables, and edited the manuscript.

Jyotika Bahuguna helped interpret the MEM model, helped improve the model implementations, and edited the manuscript.

Joshua Cape supervised the statistical analyses, helped with the interpretation, and edited the manuscript.

Bowei Ouyang helped implement the MEM models, modified the elapy code, helped interpret the MEM model, and edited the manuscript.

Satish Iyengar supervised the statistical analyses, helped with the interpretation, and edited the manuscript.

Konasale Prasad conceived the idea and the hypotheses, suggested analytical models, conducted some of the statistical analyses, edited the manuscript, and obtained the necessary funds.

## Data availability

Data are available through the National Data Archive of the NIMH

## Supplemental Materials

### Supplement 1: MRI processing and abbreviations

In image preprocessing, the T1w image was down-sampled to the fMRI images, followed by nearest-neighbor interpolation of the subject-specific atlas allowing us to map cortical boundaries to the native fMRI, obviating the need to warp or alter the fMRI data into standard space. The average voxel activation at each timepoint was then calculated for each brain region, producing a time-series of fMRI activation per individual. A full list of abbreviations related to the atlas and analysis are listed in the table below.

**Supplemental Table S1.1:** The Glasser atlas consists of 180 cortical areas per hemisphere. The areas can be grouped into well-defined networks such as DMN. Atlas abbreviations are above. The regions in the table had reduced first-order BOLD power associated with the psychosis group. Analytical abbreviations are listed below.

| Context | Abbreviation | Meaning |
| --- | --- | --- |
| Glasser atlas | DMN | Default Mode Network |
|  | RSC | Retrosplenial Complex |
|  | POS1 | Parietal-Occipital Sulcus area 1 |
|  | POS2 | Parietal-Occipital Sulcus area 2 |
|  | v23ab | Brodmann area 23 ventral subdivisions a and b |
|  | d23ab | Brodmann area 23 dorsal subdivisions a and b |
|  | 31pv | Brodmann area 31 posterior ventral |
|  | a24 | Brodmann area 24 anterior |
|  | p32 | Brodmann area 32 posterior |
|  | 10r | Brodmann area 10 rostral |
|  | TPOJ3 | Temporo-Parieto-Occipital Junction area 3 |
|  | IP1 | Intraparietal area 1 |
|  | Pfm | Parietal area F medial |
|  | PGi | Parietal area G inferior |
|  | PGs | Parietal area G superior |
|  | 31pd | Broadman area 31 posterior dorsal |
|  | 31a | Broadman area 31 anterior |
|  | s32 | Broadmann area 32 subcallosal |
|  | DAN | Dorsal Attention Network |
|  | MST | Medial Superior Temporal Area |
|  | FEF | Frontal Eye Field |
|  | 7PC | Brodmann area 7 Parietal Cortex |
|  | 7am | Brodmann area Seven Anterior-Medial |
|  | LIPd | Lateral IntraParietal area dorsal |
|  | LIPv | Lateral IntraParietal area ventral |
|  | VIP | Ventral IntraParietal complex |
|  | MT | Middle Temporal area |
|  | 6a | Brodmann area 6 subdivision a |
|  | AIP | Anterior IntraParietal area |
|  | PH | Parahippocampal area |
|  | V4t | Visual area 4 transitional |
| Analysis | MEM | Maximum Entropy Model |
|  | h | First-Order MEM Parameter |
|  | J | Second-Order MEM Parameter |
|  | BOLD | Blood Oxygen Level Dependent |
|  | FC | Functional Connectivity |
|  | DFC | Dynamic Functional Connectivity |
|  | CC | Clustering Coefficient |
|  | BC | Betweenness Centrality |
|  | AP | Affective Psychosis |
|  | NAP | Non-Affective Psychosis |
|  | LCPZE | lifetime antipsychotic exposure in chlorpromazine equivalents |

### Supplement 2: First-Order Results

The preprocessed BOLD fMRI signals from the 58 regions-of-interest, including 34 bilateral DMN and 24 bilateral DAN nodes, were compared before z-scoring by group (Figure S.2.1). There were no significant differences in any regional BOLD fMRI amplitude values within the AP fMRI acquisition, with all subjects having the same relative BOLD amplitudes. Glasser atlas regions constituting DMN and DAN are shown in the table below Figure S.2.1 where the DMN is the top row and the DAN is the bottom row, and all nodes are bilateral.

**Supplemental Figure S2.1.**
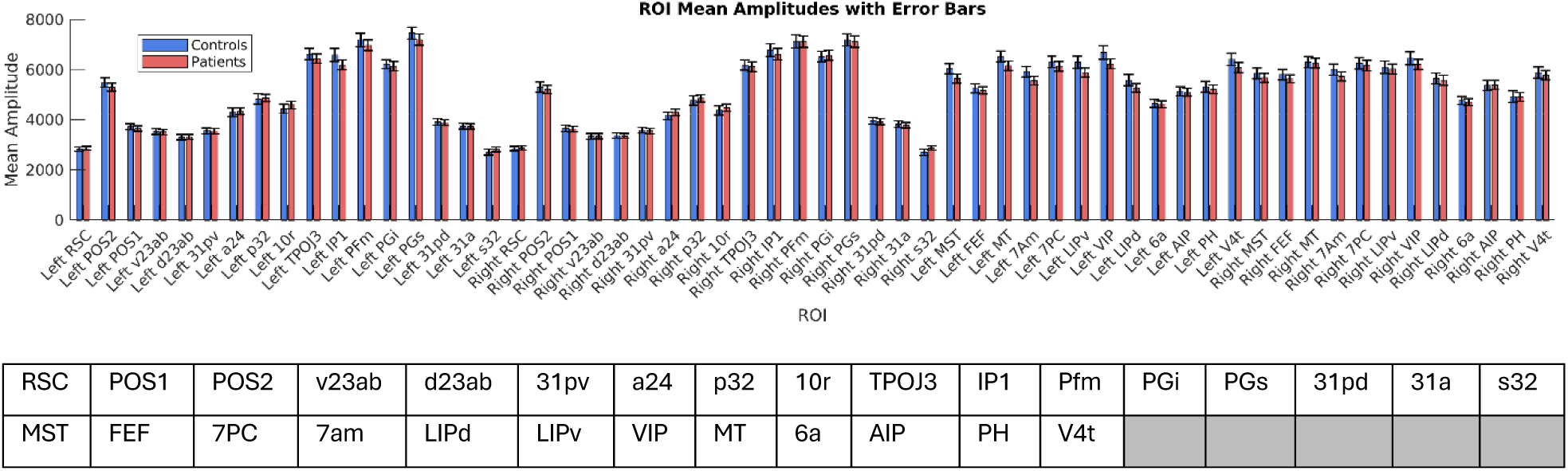
Bar Region inclusion. Bar graph on top plot shows average fMRI amplitude per region in patients (red) and controls (blue). No significant differences are observed. In the table, with the DMN nodes in the top row, and DAN in the bottom row.

The DMN and DAN were examined in the context of the whole brain for the first-order frequency band analysis only. For each participant, and each brain region (or node), the preprocessed fMRI timeseries (377 nodes x 410 timepoints) were analyzed using a Fourier transform. There were 360 cortical nodes and 23 subcortical nodes. Canonical resting-state frequency bands were then defined according to standard fMRI conventions: Slow 5 (0.01– 0.027 Hz), Slow-4 (0.027–0.073 Hz), Slow-3 (0.073–0.198 Hz), and Slow-2 (0.198–0.50 Hz). For each subject, mean band power was extracted across these four frequency ranges for every brain region. These features were restructured into a set of features representing the average power within each brain region and frequency band and combined with demographic covariates (age, sex, group, and subject ID) to generate a comprehensive feature table. The resulting dataset was exported as a .csv file for subsequent multivariate analysis of covariance (MANCOVA). All statistical analyses on whole-brain first order band power were conducted in IBM SPSS Statistics version 32. The tables below contain the results for the whole-brain, frequency-band specific, first-order analyses.

Within the DMN and DAN, 3 of the 58 regions showed significantly altered BOLD frequency power in band 5 in a MANCOVA model by including age, sex, and lifetime antipsychotic medication exposure measured as chlorpromazine equivalent (Wilk’s λ=0.52, F=1.63, p=0.016; post-hoc between-subjects’ effects, Bonferroni corrected p, all<0.05) (Figure Below). These regions were: right 31pd (posterior cingulate, EP 9.89±13.42; HC 7.76±7.07; F=4.62, p=0.03), right AIP (superior parietal, EP 7.12±8.07; HC 6.08±5.75; F=4.59, p=0.03), and left PFm (inferior parietal, EP 28.14±33.10; HC 48.30±57.48; F=4.10, p=0.045).

We observed a significant negative correlation between microstate switching and power in every band. The negative correlation was relatively weaker at faster frequency bands suggesting that power in higher frequency bands, which are associated with respiratory artifacts (slow-3 and slow-2), is less associated with microstate switching compared to slower frequency bands (slow-5, slow-4). Slower frequency bands were more strongly associated with microstate switching than slow-3 and slow-2 at the individual level (Main Text Figure 1B). In both whole-population and within-group ANOVA tests, all post-hoc band-wise differences in switching-to-power correlation were significant at p<0.001.

The top-switching network was selected under the assumption that nodes exhibiting the highest switching activity are likely to be the most informative with respect to network dynamics. However, to ensure that our findings were not solely driven by this assumption or biased toward the most active nodes, we also examined the bottom-switching network, comprising the nodes with the lowest switching activity. Analyzing both extremes of the switching spectrum allowed us to assess whether the observed patterns were specific to highly dynamic nodes or reflected more general network properties.

Among the top ten most frequently switching and bottom ten least frequently switching nodes for DMN and DAN (40 nodes), 21 showed significant microstate switching frequency differences between patients and controls (Main Text Figure 1C). Patients, on average, had more microstate switching at the group level; differences were detected in both top and bottom switching DAN and top switching DMN but not bottom switching DMN.

A MANCOVA model with all 377 brain regions for each band did not show overall model significance for BOLD signal intensity after covarying for age, sex, and lifetime exposure to antipsychotics converted to CPZ equivalents (Slow band 5: Wilk’s λ=0.04, F=1.74, p=0.55; Slow band 4: Wilk’s λ=0.001, F=4.18, p=0.38; Slow band 3: Wilk’s λ=0.004, F=1.68, p=0.56; Slow band 2: Wilk’s λ=0.0001, F=14.96, p=0.20). However, post hoc tests of between-subjects’ effects for each of these bands showed approximately 10% of the regions were significantly different; among these regions approximately 50% showed reduced BOLD signals among patients in each of these bands while the other 50% showed higher BOLD responses compared to controls. Observed regions were unique to each band with no overlap. Effect sizes were higher for reduced BOLD responses in patients (Cohen’s d ≈ 0.50) compared to controls while regions showing higher BOLD responses in patients had relatively smaller effect sizes (Cohen’s d ≈ 0.20).

### Slow 2 band

For Slow-2, the fastest frequency band, the largest effects emerged in HCP regions within auditory association and temporal cortices. While subcortical structures (globus pallidus and putamen) showed reduced BOLD signals, anterior cingulate regions primarily showed reduced BOLD signals. The dorsolateral prefrontal cortex showed both higher and lower BOLD signal intensity depending on HCP parcel.

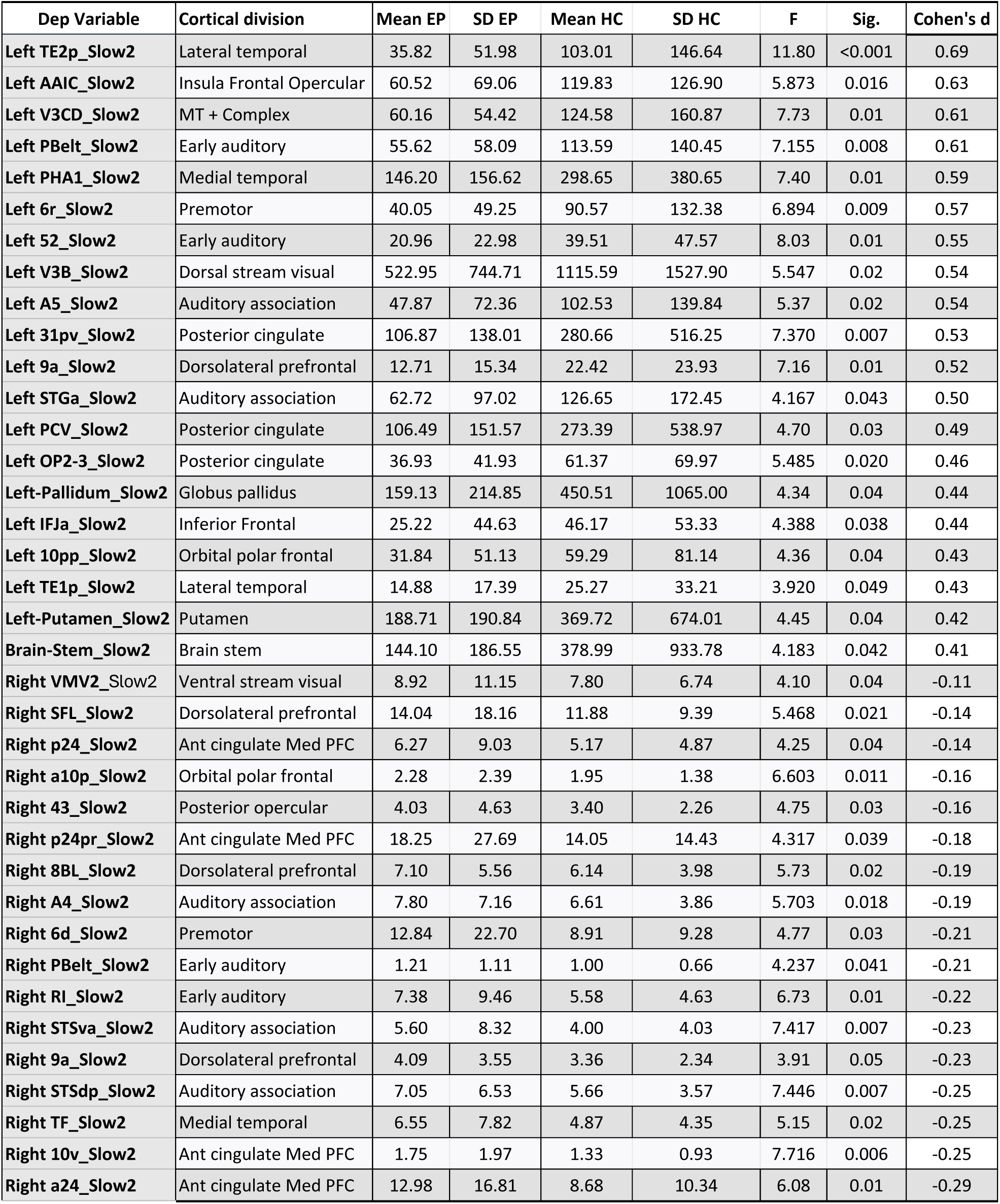

| Dep Variable | Cortical division | Mean EP | SD EP | Mean HC | SD HC | F | Sig. | Cohen's d |
| --- | --- | --- | --- | --- | --- | --- | --- | --- |
| Left TE2p_Slow2 | Lateral temporal | 35.82 | 51.98 | 103.01 | 146.64 | 11.80 | <0.001 | 0.69 |
| Left AAIC_Slow2 | Insula Frontal Opercular | 60.52 | 69.06 | 119.83 | 126.90 | 5.873 | 0.016 | 0.63 |
| Left V3CD_Slow2 | MT + Complex | 60.16 | 54.42 | 124.58 | 160.87 | 7.73 | 0.01 | 0.61 |
| Left PBelt_Slow2 | Early auditory | 55.62 | 58.09 | 113.59 | 140.45 | 7.155 | 0.008 | 0.61 |
| Left PHA1_Slow2 | Medial temporal | 146.20 | 156.62 | 298.65 | 380.65 | 7.40 | 0.01 | 0.59 |
| Left 6r_Slow2 | Premotor | 40.05 | 49.25 | 90.57 | 132.38 | 6.894 | 0.009 | 0.57 |
| Left 52_Slow2 | Early auditory | 20.96 | 22.98 | 39.51 | 47.57 | 8.03 | 0.01 | 0.55 |
| Left V3B_Slow2 | Dorsal stream visual | 522.95 | 744.71 | 1115.59 | 1527.90 | 5.547 | 0.02 | 0.54 |
| Left A5_Slow2 | Auditory association | 47.87 | 72.36 | 102.53 | 139.84 | 5.37 | 0.02 | 0.54 |
| Left 31pv_Slow2 | Posterior cingulate | 106.87 | 138.01 | 280.66 | 516.25 | 7.370 | 0.007 | 0.53 |
| Left 9a_Slow2 | Dorsolateral prefrontal | 12.71 | 15.34 | 22.42 | 23.93 | 7.16 | 0.01 | 0.52 |
| Left STGa_Slow2 | Auditory association | 62.72 | 97.02 | 126.65 | 172.45 | 4.167 | 0.043 | 0.50 |
| Left PCV_Slow2 | Posterior cingulate | 106.49 | 151.57 | 273.39 | 538.97 | 4.70 | 0.03 | 0.49 |
| Left OP2-3_Slow2 | Posterior cingulate | 36.93 | 41.93 | 61.37 | 69.97 | 5.485 | 0.020 | 0.46 |
| Left-Pallidum_Slow2 | Globus pallidus | 159.13 | 214.85 | 450.51 | 1065.00 | 4.34 | 0.04 | 0.44 |
| Left IFJa_Slow2 | Inferior Frontal | 25.22 | 44.63 | 46.17 | 53.33 | 4.388 | 0.038 | 0.44 |
| Left 10pp_Slow2 | Orbital polar frontal | 31.84 | 51.13 | 59.29 | 81.14 | 4.36 | 0.04 | 0.43 |
| Left TE1p_Slow2 | Lateral temporal | 14.88 | 17.39 | 25.27 | 33.21 | 3.920 | 0.049 | 0.43 |
| Left-Putamen_Slow2 | Putamen | 188.71 | 190.84 | 369.72 | 674.01 | 4.45 | 0.04 | 0.42 |
| Brain-Stem_Slow2 | Brain stem | 144.10 | 186.55 | 378.99 | 933.78 | 4.183 | 0.042 | 0.41 |
| Right VMV2_Slow2 | Ventral stream visual | 8.92 | 11.15 | 7.80 | 6.74 | 4.10 | 0.04 | -0.11 |
| Right SFL_Slow2 | Dorsolateral prefrontal | 14.04 | 18.16 | 11.88 | 9.39 | 5.468 | 0.021 | -0.14 |
| Right p24_Slow2 | Ant cingulate Med PFC | 6.27 | 9.03 | 5.17 | 4.87 | 4.25 | 0.04 | -0.14 |
| Right a10p_Slow2 | Orbital polar frontal | 2.28 | 2.39 | 1.95 | 1.38 | 6.603 | 0.011 | -0.16 |
| Right 43_Slow2 | Posterior opercular | 4.03 | 4.63 | 3.40 | 2.26 | 4.75 | 0.03 | -0.16 |
| Right p24pr_Slow2 | Ant cingulate Med PFC | 18.25 | 27.69 | 14.05 | 14.43 | 4.317 | 0.039 | -0.18 |
| Right 8BL_Slow2 | Dorsolateral prefrontal | 7.10 | 5.56 | 6.14 | 3.98 | 5.73 | 0.02 | -0.19 |
| Right A4_Slow2 | Auditory association | 7.80 | 7.16 | 6.61 | 3.86 | 5.703 | 0.018 | -0.19 |
| Right 6d_Slow2 | Premotor | 12.84 | 22.70 | 8.91 | 9.28 | 4.77 | 0.03 | -0.21 |
| Right PBelt_Slow2 | Early auditory | 1.21 | 1.11 | 1.00 | 0.66 | 4.237 | 0.041 | -0.21 |
| Right RI_Slow2 | Early auditory | 7.38 | 9.46 | 5.58 | 4.63 | 6.73 | 0.01 | -0.22 |
| Right STSva_Slow2 | Auditory association | 5.60 | 8.32 | 4.00 | 4.03 | 7.417 | 0.007 | -0.23 |
| Right 9a_Slow2 | Dorsolateral prefrontal | 4.09 | 3.55 | 3.36 | 2.34 | 3.91 | 0.05 | -0.23 |
| Right STSdp_Slow2 | Auditory association | 7.05 | 6.53 | 5.66 | 3.57 | 7.446 | 0.007 | -0.25 |
| Right TF_Slow2 | Medial temporal | 6.55 | 7.82 | 4.87 | 4.35 | 5.15 | 0.02 | -0.25 |
| Right 10v_Slow2 | Ant cingulate Med PFC | 1.75 | 1.97 | 1.33 | 0.93 | 7.716 | 0.006 | -0.25 |
| Right a24_Slow2 | Ant cingulate Med PFC | 12.98 | 16.81 | 8.68 | 10.34 | 6.08 | 0.01 | -0.29 |

### Slow 3 band

For the slow-3 band, the largest effects emerged in HCP regions within dorsolateral prefrontal cortex and inferior frontal cortex, as well as some temporal lobe regions. Other temporal lobe structures and the parietal-temporal junction area (MT+ complex)) showed reduced BOLD signals among patients. Early auditory cortex in particular showed both higher and lower BOLD signal intensity depending on HCP parcel.

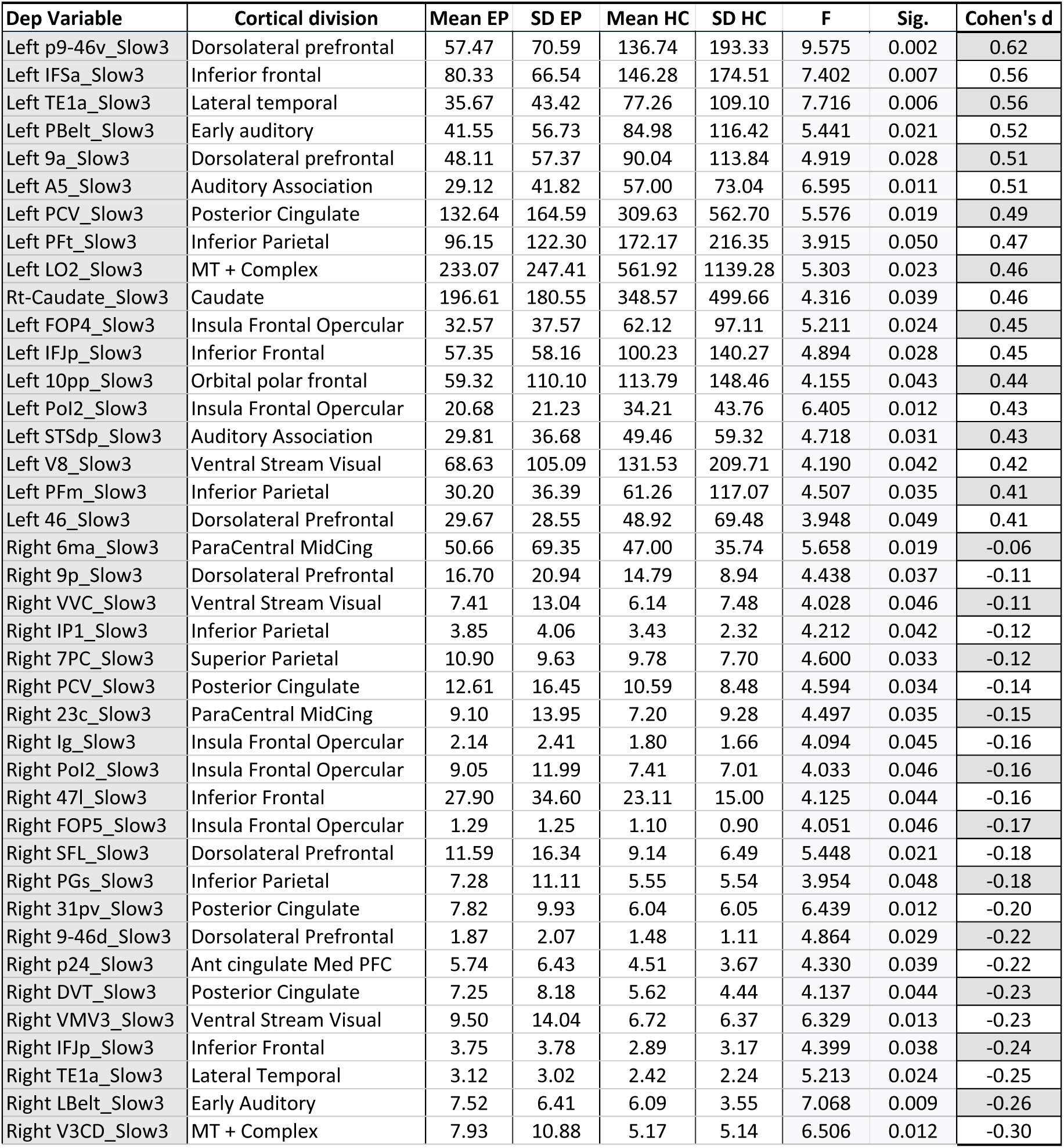

### Slow 4 band

For the slow-4 band, the largest effects emerged in HCP regions within temporal lob structure including the early auditory cortex and auditory association cortex. While subcortical structures (caudate and thalamus) also showed increased slow-4 power as compared to controls. Premotor, cingulate, dorsolateral prefrontal, and some auditory cortex areas show decreased slow-4 power compared to controls.

| Dep Variable | Cortical division | Mean EP | SD EP | Mean Hc | SD HC | F | Sig. | Cohen's d |
| --- | --- | --- | --- | --- | --- | --- | --- | --- |
| Left PBelt_Slow4 | Early Auditory | 49.74 | 53.79 | 102.84 | 118.32 | 9.809 | 0.002 | 0.64 |
| Left AIP_Slow4 | Superior Parietal | 76.49 | 71.51 | 149.99 | 187.43 | 7.985 | 0.005 | 0.59 |
| Left IFJa_Slow4 | Inferior Frontal | 57.86 | 54.56 | 117.24 | 153.61 | 7.455 | 0.007 | 0.59 |
| Left STSda_Slow4 | Auditory Association | 86.97 | 72.71 | 148.57 | 151.71 | 4.376 | 0.038 | 0.57 |
| Left 52_Slow4 | Early Auditory | 23.70 | 26.43 | 45.65 | 57.39 | 8.083 | 0.005 | 0.55 |
| Left 6r_Slow4 | Premotor | 60.10 | 82.64 | 128.20 | 178.83 | 7.173 | 0.008 | 0.54 |
| Left-Thalamus_Slow4 | Thalamus | 132.25 | 219.70 | 304.28 | 475.52 | 4.396 | 0.038 | 0.52 |
| Right-Caudate_Slow4 | Caudate | 376.59 | 530.05 | 733.88 | 1015.35 | 3.987 | 0.048 | 0.48 |
| Left AAIC_Slow4 | Insula Frontal Opercular | 38.75 | 52.17 | 72.00 | 94.54 | 6.191 | 0.014 | 0.47 |
| Left-Caudate_Slow4 | Caudate | 346.41 | 488.37 | 743.56 | 1270.64 | 4.830 | 0.029 | 0.47 |
| Left-Putamen_Slow4 | Putamen | 168.75 | 150.93 | 358.80 | 660.07 | 5.147 | 0.025 | 0.46 |
| Left A5_Slow4 | Auditory Association | 48.03 | 52.19 | 90.68 | 144.89 | 4.183 | 0.042 | 0.44 |
| Left PeEc_Slow4 | Medial temporal | 43.88 | 49.57 | 76.77 | 107.38 | 4.775 | 0.030 | 0.44 |
| Left s32_Slow4 | Ant cingulate Med PFC | 17.97 | 22.46 | 28.70 | 30.07 | 3.886 | 0.050 | 0.42 |
| Right PBelt_Slow4 | Early Auditory | 5.63 | 6.73 | 5.02 | 5.80 | 4.821 | 0.030 | -0.10 |
| Right TPOJ3_Slow4 | TPO | 26.31 | 33.95 | 23.31 | 20.51 | 6.692 | 0.011 | -0.10 |
| Right s6-8_Slow4 | Dorsolateral prefrontal | 4.08 | 3.40 | 3.69 | 2.62 | 4.126 | 0.044 | -0.12 |
| Right TGd_Slow4 | Lateral temporal | 4.79 | 4.63 | 4.27 | 2.72 | 4.053 | 0.046 | -0.13 |
| Right V4t_Slow4 | MT + Complex | 10.64 | 14.34 | 8.79 | 8.47 | 4.240 | 0.041 | -0.15 |
| Right 5L_Slow4 | ParaCental MidCing | 11.66 | 20.29 | 8.99 | 7.92 | 4.705 | 0.032 | -0.16 |
| Right MBelt_Slow4 | Early Auditory | 11.09 | 13.78 | 9.04 | 7.13 | 4.549 | 0.034 | -0.17 |
| Right 10pp_Slow4 | Orbital polar frontal | 0.86 | 0.74 | 0.73 | 0.65 | 4.381 | 0.038 | -0.17 |
| Right 25_Slow4 | Ant cingulate Med PFC | 10.66 | 11.13 | 8.86 | 7.72 | 4.452 | 0.036 | -0.18 |
| Right p9-46v_Slow4 | Dorsolateral prefrontal | 9.73 | 12.07 | 7.68 | 6.63 | 5.470 | 0.021 | -0.20 |
| Right 31pv_Slow4 | Posterior cingulate | 6.00 | 8.41 | 4.35 | 4.80 | 6.430 | 0.012 | -0.23 |
| Right FOP5_Slow4 | Insula Frontal Opercular | 2.02 | 2.00 | 1.60 | 1.47 | 4.463 | 0.036 | -0.23 |
| Right TE1a_Slow4 | Lateral temporal | 5.41 | 6.59 | 4.05 | 4.43 | 6.364 | 0.013 | -0.23 |
| Right a9-46v_Slow4 | Dorsolateral prefrontal | 1.79 | 2.51 | 1.24 | 1.16 | 5.164 | 0.024 | -0.26 |
| Right OP1_Slow4 | Posterior opercular | 6.56 | 10.22 | 4.39 | 3.06 | 3.914 | 0.050 | -0.26 |
| Right 46_Slow4 | Dorsolateral prefrontal | 11.45 | 12.73 | 8.55 | 7.68 | 4.756 | 0.031 | -0.26 |
| Right LBelt_Slow4 | Early auditory | 5.87 | 4.76 | 4.60 | 3.36 | 6.792 | 0.010 | -0.30 |
| Right a24_Slow4 | Ant cingulate Med PFC | 9.89 | 13.20 | 6.43 | 7.89 | 7.460 | 0.007 | -0.30 |
| Left 6v_Slow4 | Premotor | 16.52 | 26.61 | 9.77 | 10.10 | 4.561 | 0.034 | -0.30 |

### Slow 5 band

For slow-5, the slowest frequency band, the largest effects emerged in HCP regions within inferior frontal, cingulate, and again auditory association and temporal cortices. Lateral temporal structures showed the greatest reduction in BOLD slow-5 power compared to controls. The cingulate cortex showed both higher and lower BOLD signal intensity depending on HCP parcel.

**Tables S1.**
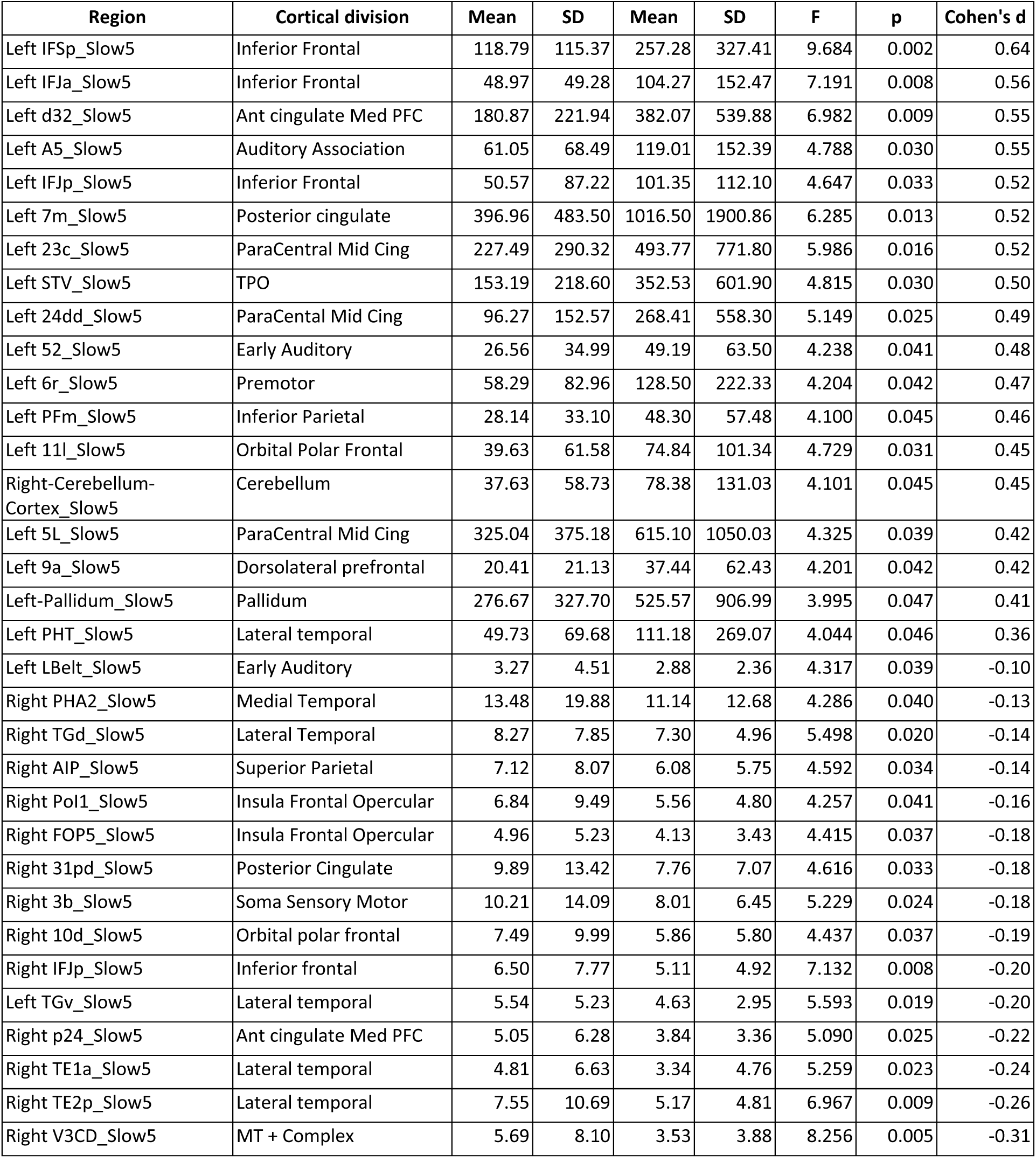
All significant results from brain-wide, group-wise, power band analysis.

### Supplement 3: Second-Order Results

At the whole-graph level, a MANCOVA model controlling for age and sex showed significant group effect across the combined global network measures (average clustering, global efficiency, modularity, and average BC; Wilks’ λ=0.94, F(3,159)=3.15, p=0.02). Post-hoc tests of between-subjects effects with Bonferroni correction showed significant group differences for all graph measures (p<0.015). Network connectivity metrics for the entire 58-node DMN+DAN network, as well as the 34-node DMN and 24-node DAN separately. Patients had lower average CC and efficiency, but higher modularity and average BC compared to controls. We observed similar results for networks built with each frequency band, except for fewer differences in slow-2 band than other bands.

At the whole-graph level, a MANCOVA controlling for age and sex revealed a significant group effect across the combined global network measures (average clustering, global efficiency, modularity, and average betweenness centrality; Wilks’ λ = 0.94, F(3,159) = 3.15, p = 0.020). Post-hoc between-subjects tests with Bonferroni correction indicated significant group differences for all measures (p < 0.015). Network connectivity metrics for the entire 58-node DMN+DAN network, as well as the 34-node DMN and 24-node DAN separately, are presented in Table 2. Patients had lower average clustering and efficiency, but higher modularity and average betweenness centrality compared to controls.

|  |  | DMN |  | DAN |  | DMN+DAN |  |
| --- | --- | --- | --- | --- | --- | --- | --- |
| Metric | Group | Mean | SD | Mean | SD | Mean | SD |
| Clustering | Controls | **0.41 | 0.09 | **0.49 | 0.14 | **0.36 | 0.11 |
|  | Patients | **0.36 | 0.11 | **0.43 | 0.13 | **0.30 | 0.11 |
| Modularity | Controls | **0.07 | 0.03 | 0.06 | 0.04 | **0.12 | 0.05 |
|  | Patients | **0.09 | 0.03 | 0.07 | 0.04 | **0.15 | 0.06 |
| Efficiency | Controls | **0.47 | 0.08 | **0.53 | 0.11 | **0.43 | 0.09 |
|  | Patients | **0.42 | 0.09 | **0.48 | 0.11 | **0.39 | 0.09 |
| Betweenness | Controls | 23.37 | 11.12 | 12.02 | 9.36 | **54.35 | 27.54 |
|  | Patients | 26.61 | 12.57 | 14.03 | 7.80 | **67.59 | 28.94 |

Graph metrics for second-order comparisons. Asterisks indicate significance at \*\* $p<0.01$ .

At the individual-edge level the differences between groups are visually noticeable: the average FC for the DMN+DAN is shown by group in **Figures 2A and B**. Of the 1,653 unique edges in this 58-node network, 181 of them, or about 10%, demonstrated different edge weight distributions between groups at the alpha=0.001 level in KS-tests, with more negatively weighted edges in patients (**Supplemental Section S.2**). Overall, this shows that patients have weaker connections at the edge-level, corroborated by lower connection strength, lower clustering, and higher modularity in psychosis patients at the node and graph levels.

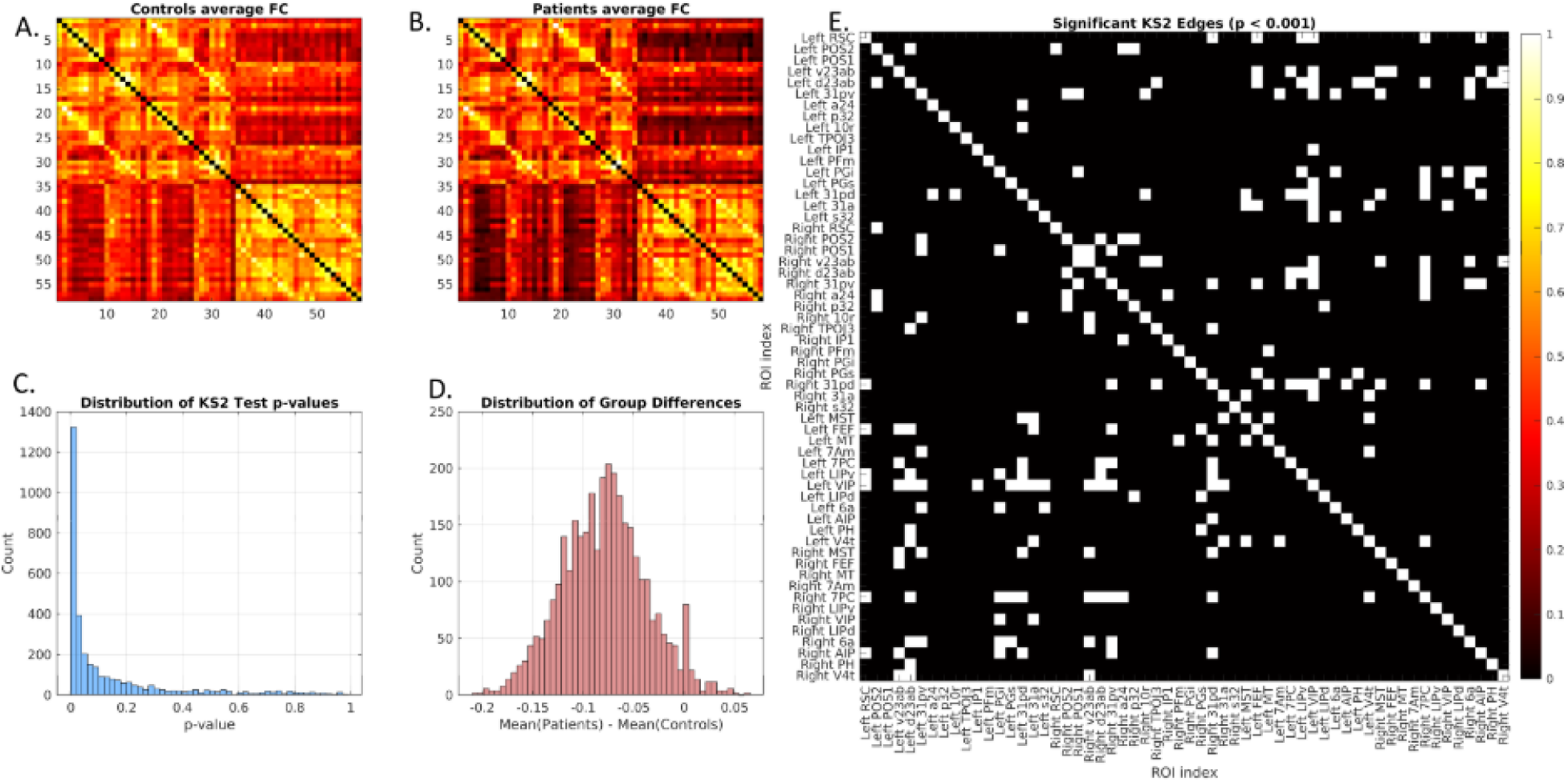
Second Order Results. **A)** The control-group average FC for edges between all DMN and DAN nodes. **B)** The patient-group average FC for all unique DMN and DAN node-pairs. **C)** Histogram of p-values for two-sample KS-test of FC edge weight difference by group over all tests. **D)** Histogram of difference in group level average edge weight. **E)** The binary outcome of null hypothesis rejection at level alpha 0.001 for the two-sample KS-test by edge.

The four canonical frequency bands described above (slow-5, slow-4, slow-3, and slow-2) were applied as bandpass filters to the preprocessed timeseries frequency domain data. Then, the filtered frequency domains were inverse Fourier transformed back into the time domain, creating four band-filtered timeseries per region and per subject. Next, the functional connectomes were created for the 58-node DMN+DAN for each frequency band. The same graph measures were then calculated for the frequency-band specific functional connectomes as were calculated for the full-spectrum FC shown in the main text. Groupwise (patient-control) differences were then examined using two-sample t-test for each measure.

These analyses were repeated for networks constructed separately for each frequency band, yielding similar results, but with fewer differences in slow-2 than the other frequency bands (Supplemental Section S.2). Based on these findings, subsequent analyses focused on full-spectrum functional connectivity. At the nodal level, a separate MANCOVA controlling for age and sex did not reveal a significant multivariate effect of group on the combined nodal network measures (CC, BC, and strength; Wilks’ λ=0.99, F(2,160)=0.05, p=0.95). However, Bonferroni-corrected tests of between-subjects effects showed higher BC (p=0.011) but lower CC (p=0.015) and strength (p=0.0079) in patients compared to controls.

At the individual-edge level, the differences between groups were noticeable in the average FC for the DMN+DAN (Supplemental Figures 2A and B). Of the 1,653 unique edges in this 58-node network, 181 of them (≈10%), demonstrated different edge weight distributions between groups at the α=0.001 in KS-tests, with more negatively weighted edges in patients (Supplemental Information 2). Overall, this shows that patients have weaker connections at the edge level, corroborated by lower connection strength, lower clustering, and higher modularity at the node and graph levels.

About 10% of the edges demonstrated different edge weight distributions between groups at the α=0.001 level in KS-tests, with more negatively weighted edges in patients, These observations suggest that patients have weaker connections at the edge level, corroborated by lower connection strength, lower clustering, and higher modularity at the node and graph levels (Supplemental Figure 3). Of the 816 unique edges that represent the direct connections between DMN and DAN, at the individual level, there were more negatively weighted edges in the patient group (on average 129 per patient) than the control group (approximately 74 per person). Among patients, the average minimum edge weight for edges between DMN and DAN was r =–0.39 (±0.195), compared to r= –0.21 (±0.192) among controls. A two-sample KS-test comparing the FC edge-weight distributions between patients and controls indicated patients have significantly more negative correlations (main text Figure 2D,E; KS =.324, p<.001), although the standard deviations were very large in both groups (patients: 196±140 negatively weighted edges; controls: 104±92).

Regarding graph metrics, a summary of the relevant findings from the main text goes as follows: Patients showed lower average clustering coefficient in the DMN+DAN(t = -3.34, p = .0011), DMN (t = -3.34, p = .0010), and DAN (t = -2.80, p = .0057), and the same findings were observed for efficiency: DMN+DAN(t = -3.33, p = .0011), DMN (t = -3.27, p = .0013), and DAN (t = -2.82, p = .0054). Modularity coefficient was significantly higher among patients in the DMN+DAN (t = 2.83, p=.0052) and the DMN (t=3.133, p=.0020), but not in the DAN. Average betweenness centrality was only significantly different in the DMN+DAN network (t = 2.83, p = .0053), but group differences were not found in either DMN or DAN alone for average betweenness centrality.

Group-level average FC networks for each frequency band are shown in Supplemental Figure 1. Visibly, the FC correlations are weaker across the board for both patients and controls at higher frequencies. A table of graph measure difference is provided in Supplemental Table 2. The results are very consistent with full spectrum FC, and all significant differences are in the same direction as the full spectrum FC. Some graph properties can be localized to specific bands.

**Supplemental Figure 1.**
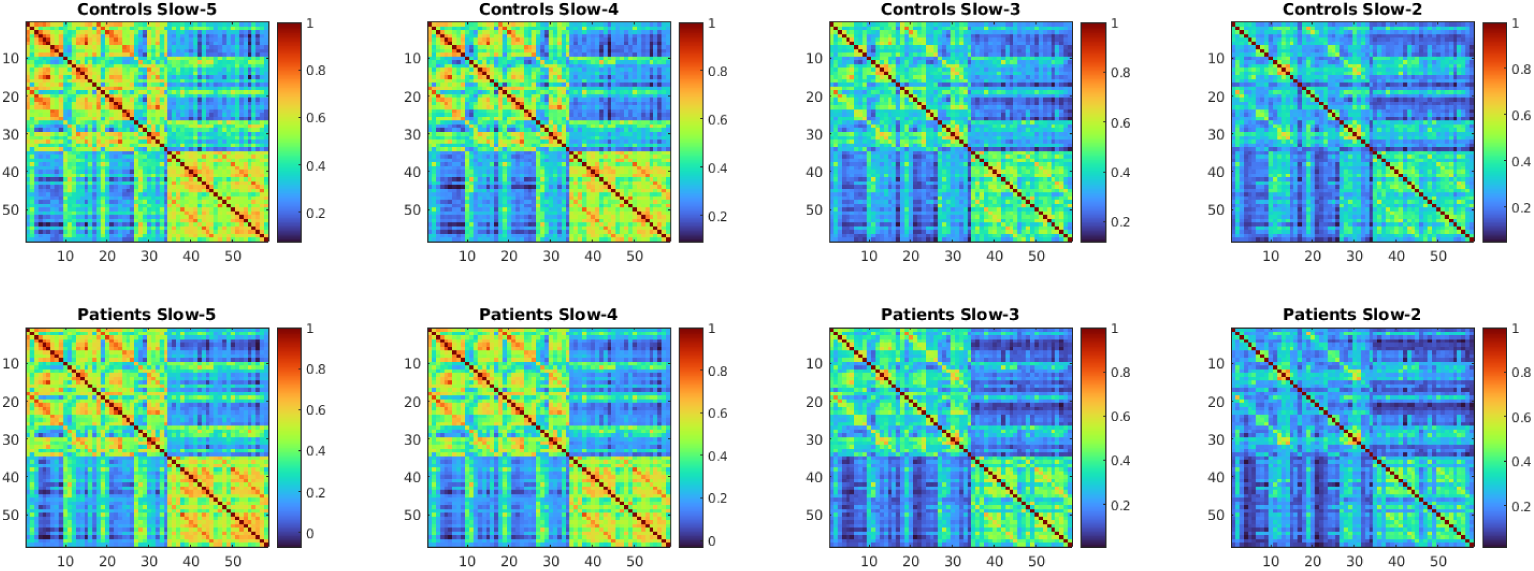
The frequency-band-specific FC group level averages. Top row: controls; bottom row: patients. From left to right: slow-5, slow-4, slow-3, slow-2. Nodal order is the same as in the main manuscript (DMN is upper left block, DAN is lower right). Inter-network connections are in the off-diagonal blocks.

**Supplemental Table S2.** DMN+DAN. The frequency-band-specific FC graph metric comparison results for DMN+DAN (58 nodes).

|  | Measure | Mean Patients | SD Patients | Mean Controls | SD Controls | T value | p Value |
| --- | --- | --- | --- | --- | --- | --- | --- |
| Slow-5 | Clustering | 0.42 | 0.111 | 0.48 | 0.129 | *-3.03 | 0.0029 |
|  | Modularity | 0.18 | 0.424 | 0.13 | 0.049 | 1.88 | 0.0618 |
|  | Efficiency | 0.52 | 0.080 | 0.56 | 0.091 | *-2.99 | 0.0032 |
|  | Betweenness | 46.73 | 19.170 | 34.08 | 18.046 | *4.01 | 9.3391e-05 |
| Slow-4 | Clustering | 0.37 | 0.128 | 0.42 | 0.125 | *-2.31 | 0.0222 |
|  | Modularity | 0.17 | 0.061 | 0.15 | 0.059 | *2.07 | 0.0396 |
|  | Efficiency | 0.45 | 0.093 | 0.49 | 0.090143 | *-2.99 | 0.0230 |
|  | Betweenness | 60.27 | 30.818 | 45.17 | 25.519 | *4.01 | 0.0019 |
| Slow-3 | Clustering | 0.29 | 0.098 | 0.37 | 0.105 | *-4.75 | 4.4887e-06 |
|  | Modularity | 0.17 | 0.051 | 0.13 | 0.049 | *4.40 | 1.9336e-05 |
|  | Efficiency | 0.36 | 0.077 | 0.42 | 0.080 | *-4.55 | 1.0568e-05 |
|  | Betweenness | 76.13 | 35.009 | 58.78 | 29.048 | *3.19 | 0.0017 |
| Slow-2 | Clustering | 0.26 | 0.101 | 0.30 | 0.109 | *-2.37 | 0.0191 |
|  | Modularity | 0.14 | 0.043 | 0.13 | 0.049 | 1.26 | 0.2081 |
|  | Efficiency | 0.33 | 0.087 | 0.36 | 0.089 | *-2.37 | 0.0187 |
|  | Betweenness | 39.73 | 24.926 | 44.72 | 26.924 | -1.18 | 0.2380 |

**Supplemental Table S2.** DMN. The frequency-band-specific FC graph metric comparison results for DMN only (34 nodes).

|  | Measure | Mean Patients | SD Patients | Mean Controls | SD Controls | T value | p Value |
| --- | --- | --- | --- | --- | --- | --- | --- |
| Slow-5 | Clustering | 0.50 | 0.131 | 0.56 | 0.130 | *-3.02 | 0.0030 |
|  | Modularity | 0.12 | 0.043 | 0.10 | 0.044 | *2.63 | 0.0094 |
|  | Efficiency | 0.56 | 0.096 | 0.61 | 0.095 | *-3.06 | 0.0029 |
|  | Betweenness | 19.60 | 10.804 | 14.58 | 9.421 | *2.94 | 0.0037 |
| Slow-4 | Clustering | 0.47 | 0.127 | 0.50 | 0.110 | -1.44 | 0.1508 |
|  | Modularity | 0.11 | 0.041 | 0.10 | 0.033 | 1.17 | 0.2454 |
|  | Efficiency | 0.514 | 0.096 | 0.53 | 0.080 | -1.35 | 0.1775 |
|  | Betweenness | 20.52 | 10.933 | 17.64 | 9.046 | 1.63 | 0.0925 |
| Slow-3 | Clustering | 0.36 | 0.107 | 0.43 | 0.099 | *-4.03 | 8.5547e-05 |
|  | Modularity | 0.13 | 0.043 | 0.106 | 0.038 | *3.56 | 0.0005 |
|  | Efficiency | 0.40 | 0.085 | 0.45 | 0.077 | *-3.947 | 0.0001 |
|  | Betweenness | 28.72 | 13.697 | 24.75 | 12.643 | 1.81 | 0.0724 |
| Slow-2 | Clustering | 0.28 | 0.112 | 0.32 | 0.096 | *-2.31 | 0.0219 |
|  | Modularity | 0.15 | 0.052 | 0.14 | 0.044 | *2.54 | 0.0121 |
|  | Efficiency | 0.32 | 0.94 | 0.36 | 0.079 | *-2.33 | 0.0210 |
|  | Betweenness | 14.73 | 12.652 | 18.69 | 10.510 | *-2.01 | 0.0461 |

**Supplement Table S2.**
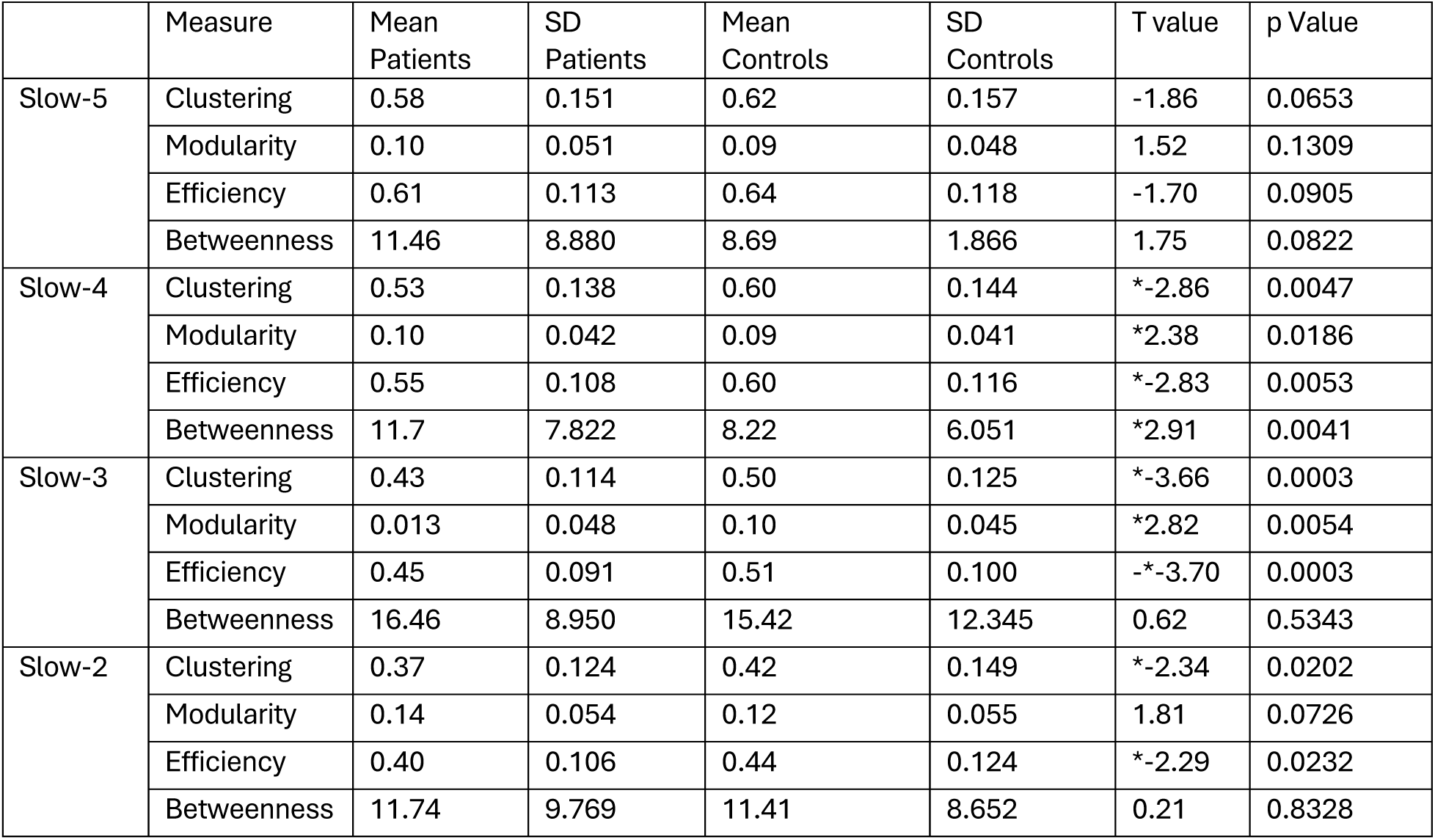
DAN. The frequency-band-specific FC graph metric comparison results for DAN only (24 nodes).

### Supplement 4: MEM methods and results

MEMs were fit using the maximum likelihood estimation (MLE). The MLE is an iterative method that runs until convergence and determines values of J and h, the interaction coefficient, and the activation bias, respectively, for each system. Once fit, the theoretical probability of a brain state can be calculated from the model parameters, J and h. This theoretical probability is defined for each unique brain state and is the expected rate of occurrence (in the whole population) of that state. Correlating the theoretical probability to the actual observed (sometimes called empirical) probability of the states demonstrates how well the MLE algorithm performed, i.e. how well the MEM explains the data. As shown in Supplemental Figure 2, all models had excellent fits (r > .99, p < .001).

**Supplemental Figure 2.**
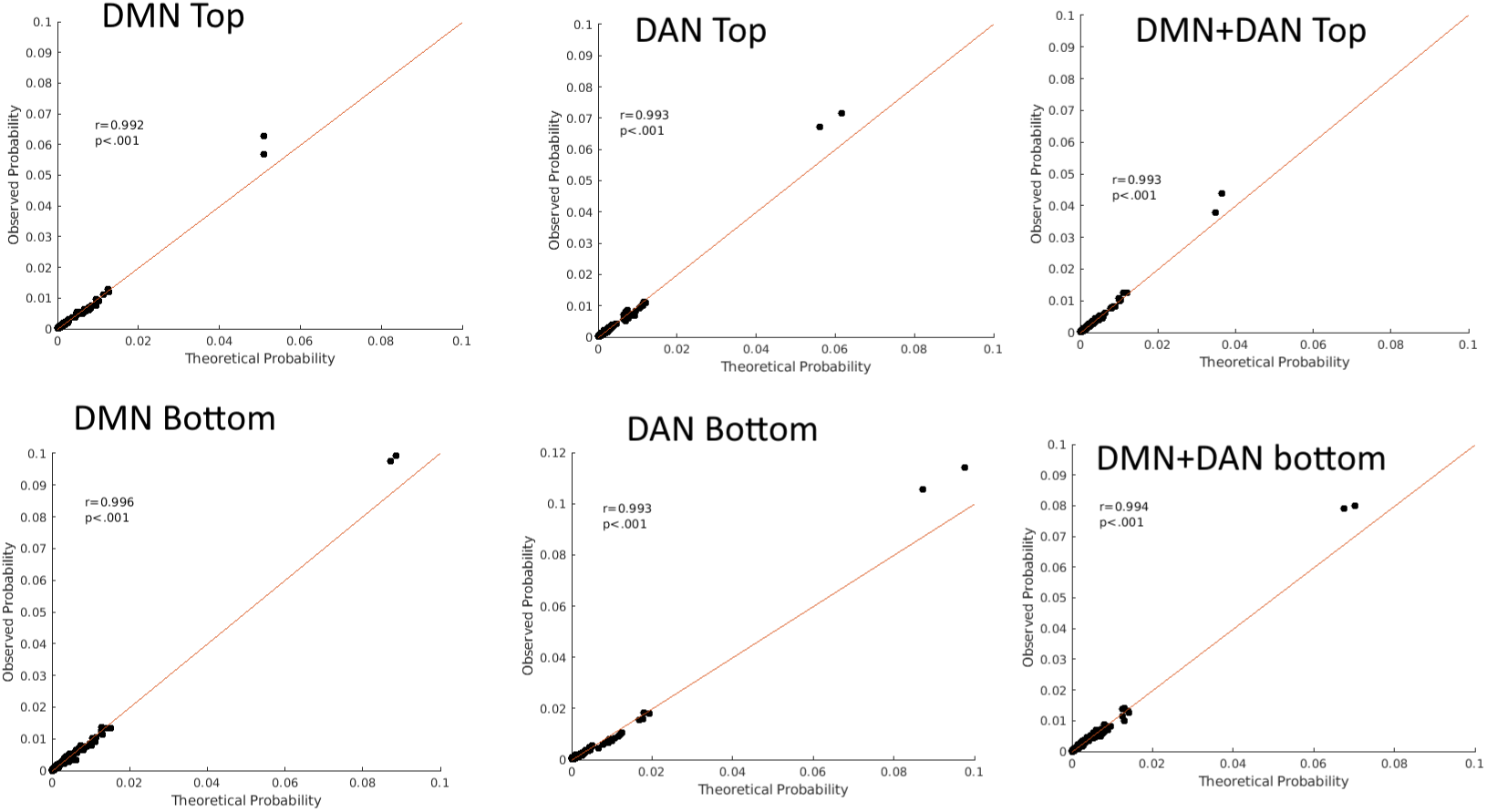
Scatter plots for whole-population MEM fit results. The x-axis shows the theoretical probability, the y-axis the observed probability. Scatter points are unique brain states. The red line is the linear least-squared squared line. All correlations are highly significant (r >. 99, p < .001).

To explain further, MEM parameters were estimated using a two-stage procedure applied to binary state vectors concatenated across all subjects and timepoints (165 subjects*410 timepoints=67,650 observations). In the first stage, pseudo-likelihood estimation was used to obtain initial parameter values via iterative coordinate ascent (learning rate α=0.5; maximum 1,000 iterations; early stopping when first-and second-order moment conditions were simultaneously satisfied within numerical tolerance). The interaction matrix J was enforced to have zero diagonal throughout. In the second stage, these pseudo-likelihood estimates served as warm-start initialization for exact maximum likelihood estimation via L-BFGS-B optimization(1), minimizing the sum of squared discrepancies between model-predicted and observed first-order activation means and second-order co-activation correlations across all 2^n^ possible states (maximum 5,000 iterations; function-value convergence tolerance ftol=1e-6). J was parameterized via its upper triangular entries and symmetrized at reconstruction to enforce symmetry. No explicit regularization penalty was applied: each 10-node MEM has 55 free parameters estimated from 67,650 observations (ratio ∼1:1,230), providing sufficient implicit regularization against extreme parameter values. The robustness of fitted parameters to node selection was confirmed by permutation analyses across 500 randomly selected 10-node subsets(1). Because parameters were estimated from a single population-level model fit to state vectors pooled across all subjects rather than separately per individual or subsample, the resulting energy landscape reflects the shared statistical structure of the full sample rather than being disproportionately shaped by any individual or subgroup, mitigating sample bias. Correlated inputs are an inherent feature of fMRI data and of the Ising framework itself, since J is explicitly designed to capture pairwise co-activation; the symmetric upper-triangular parameterization constrains each pairwise relationship to a single parameter, and initializing J from the empirical co-activation matrix, which is well-conditioned at this sample size, prevents the optimizer from beginning in regions of parameter space associated with extreme interaction weights.

We examined differences in MEM metrics among non-affective psychosis, affective psychosis patients, and controls using MANCOVA that showed a trend toward significance (Wilk’s λ=0.72, F=1.42, p=0.051). Post-hoc between-subjects’ effects for MEM metrics with Bonferroni corrections showed a similar pattern of significant differences in MEM metrics as psychosis-control comparison except for bottom distinct states showing a trend toward significance (p=0.08). Another post-hoc test to compare NAP, AP, and controls, showed that all MEM metrics with significant differences in the between-subjects’ effects tests were different between controls and NAP but not between controls and AP except top energy trace, which was the highest in NAP, intermediate in AP and the lowest in controls after Games-Howell corrections for multiple testing.

Between NAP and AP patients, no differences in the distributions of distinct states were found in any of the six systems. Between NAP and controls, differences were seen in every system (DMN-top: KS=0.33, p=0.001; DMN-bottom: KS=0.31, p=0.002; DAN-top: KS=0.30, p=0.003; DAN-bottom: KS=0.33, p-0.001; DMN+DAN-top: KS=0.34, p<0.001; DMN+DAN-bottom: KS=0.33, p<0.001). Between AP and controls, significant differences were observed in three systems (DMN-bottom: KS=0.32,p=0.037; DAN-top: KS=0.37, p=0.012; DMN-DAN-top: KS=0.448, p<0.001).

**Supplemental Figure 3:**
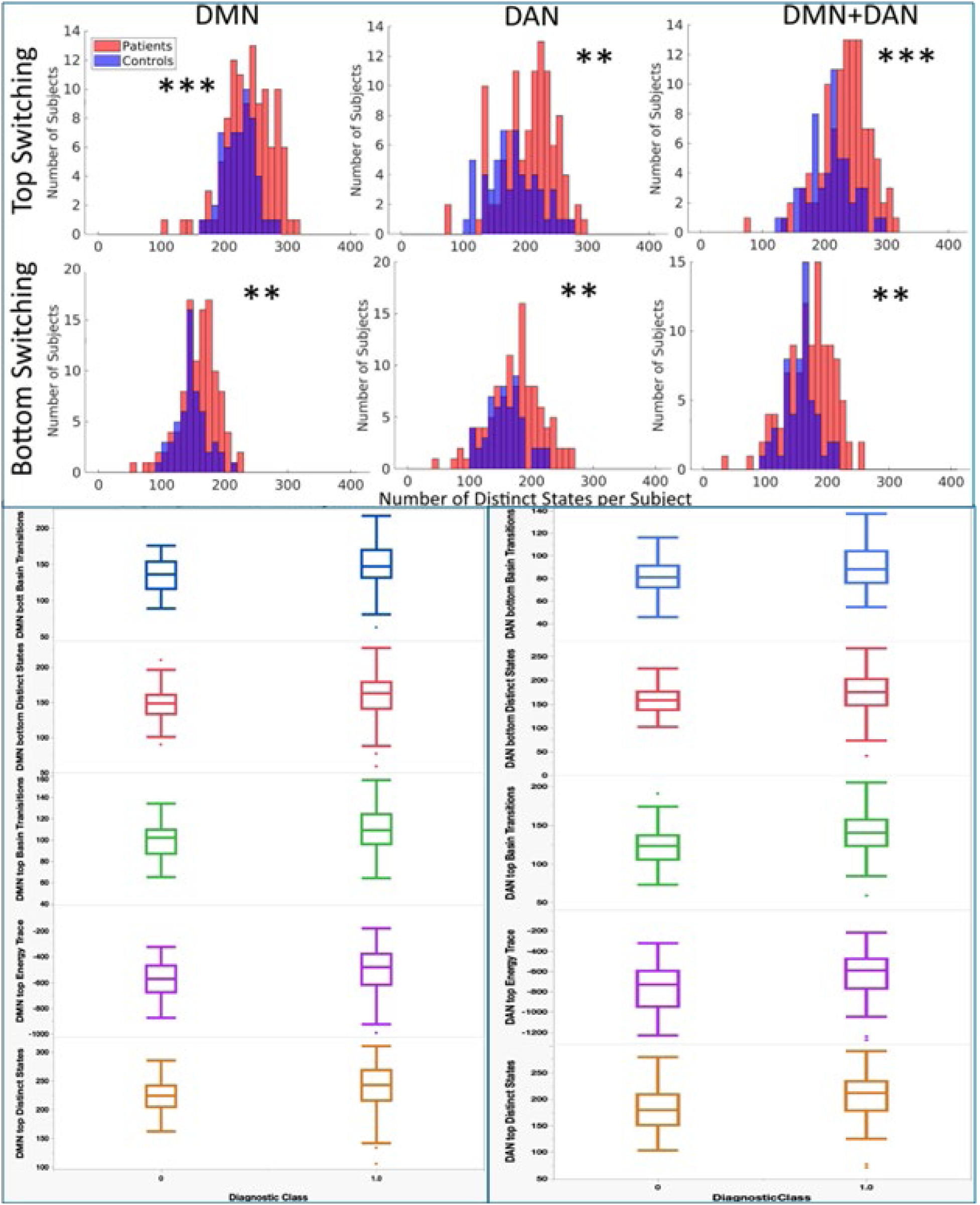
Top: The total number of distinct states per subject is shown in two histograms (one per group) for the six systems. Asterisks indicate significance of difference of distribution by 2-sample KS test (**p<0.01; ***p<0.001). Bottom: Differences in MEM metrics for DMN and DAN observed using MANCOVA model. Observed significance are corrected for multiple tests using Bonferroni approach.

Permutation tests were conducted for the DMN, DAN, and DMN+DAN. In each case, the selection of nodes was being permuted over 500 instances. In each instance, for a given case (say, DMN), 10 nodes are randomly selected from the system. Next, this node set (permutation) is fit using the same MEM approach described in Section 2.4 of the main text.

For each permutation, the MEM “summary variables” assessed in the main text, namely number of distinct states, energy trace, and number of basin transitions in the energy landscape are calculated for each participant. Then, group-level differences in the distribution of each variable are assessed by two-sample KS-test. This provides a distribution of over 500 permutations of the group level differences of the summary variables, for each system (Figures Below).

Results are summarized as follows: for the DMN, patients had more distinct states in 99% of permutations, higher energy trace in 96%, and more basin transitions in 96%. For the DAN, 96% of permutation had more distinct states, 65% had higher energy trace, and 77% of permutations had significantly more basin transitions in patients. For the combined DMN+DAN systems, across permutations, 99% show more distinct states for patients, 95% higher energy trace, and 96% had more basin transitions in the patient group. This corresponds to the results seen in the main text where for patients, all systems had more unique states (and permutations are roughly 96-99% likely to demonstrate this property), some systems had higher energy traces for patients (and only 65% of DAN permutations demonstrate this property), and transitions difference fall in-between (roughly 77-96%).

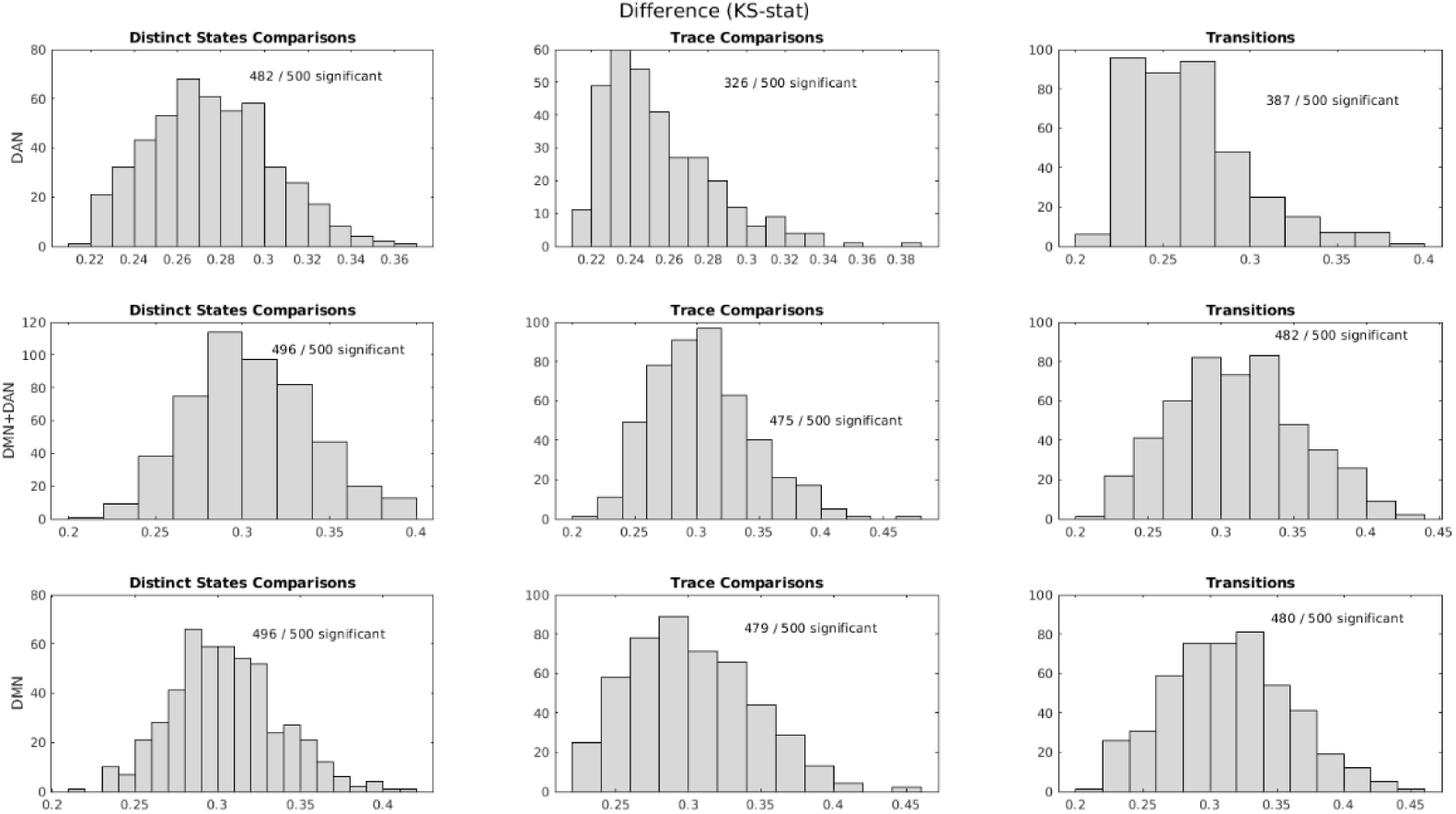
Permutation Figure (KS-statistic). In the above histograms, across the nine subplots, the rows represent the systems, which are, from top to bottom: DAN, DMN+DAN, and DMN. Column represent summary measures, which are, from left to right: distinct states, trace, and transitions. Histograms represent 500 permutations, showing the distribution of KS-statistics.

### Supplement 5: Modifications made to elapy

Modifications included: (1) replacing the iterative parameter estimation in fit_exact with L-BFGS-B optimization initialized from pseudo-likelihood estimates; (2) enforcing zero diagonal and small random initialization in fit_approx to improve convergence; (3) relabeling energy basins by the state index of their local minimum to ensure consistent cross-condition comparability; and (4) correcting the bottleneck-energy computation in the disconnectivity graph algorithm to conform to the minimax-path definition.

### Supplement 6: Correlation of MEM metrics with cognition

To formally assess whether MEM-derived metrics provide incremental predictive value for working memory performance beyond conventional first-and second-order measures, we conducted a hierarchical regression analysis. The MEM energy function takes the form E(σ) = −Σᵢhᵢσᵢ − Σᵢ < ⱼJᵢⱼσᵢσⱼ, where the h terms encode first-order regional activation and the J terms encode second-order pairwise correlation structure. Because MEM takes first-and second-order statistics as inputs and transforms them into an energy landscape from which higher-order dynamic metrics are derived, we did not treat MEM as a sequentially independent third block in a standard hierarchical regression. Instead, we compared two complementary models predicting LSWMT (N=155 with complete cognitive data). Model A entered covariates (age, sex, LCPZE, NAP/AP) in Block 1 followed by a first-order band power composite (PC1 of left IP1, left PH, right POS2 slow-5 power; 79% within-domain variance) in Block 2 and a second-order FC graph metric composite (PC1 of DMN+DAN clustering, efficiency, modularity, and betweenness centrality; 88% within-domain variance) in Block 3. Model B entered the same covariates in Block 1 followed by a MEM composite (PC1 of the five basin-transition and energy-trace metrics significantly associated with working memory; 66% within-domain variance) in Block 2. PCA composites were used within each domain because the raw FC graph metrics were collinear (r=0.99 between clustering and efficiency), which rendered individual variable entry statistically intractable (VIF>100). As a sensitivity check, we also tested Model C, which entered covariates, then MEM, then first-order, then second-order FC sequentially.

### Supplement 7: Sample description

The sample consisted of 165 subjects (109 patients and 56 controls) after excluding nine for either not having a complete cortical parcellation, failing a step of preprocessing, or not having the expected number of time points. Patients were younger than controls (t=3.33, p=0.001) with no difference in sex distribution (χ^2^=0.21, p=0.64). Approximately 76% had non-affective psychoses (NAP) and 24% had affective psychosis (AP). See the table below for demographic and clinical characteristics.

**Supplemental Table S1.1.** Clinical and demographic data. (some participants may have more than one diagnosis as these are lifetimes rates)

| Group | N | Mean Age (years) | Sd Age (years) | Females | Males | AP | NAP |
| --- | --- | --- | --- | --- | --- | --- | --- |
| Controls | 56 | 24.9 | 4.0 | 19 (34%) | 37 (66%) | 0 | 0 |
| Patients | 109 | 22.8 | 3.8 | 41 (38%) | 68 (62%) | 26 | 83 |
| Schizophrenia | 58 | 24.93 | 4.00 | 19 | 39 |  |  |
| Schizoaffective d/o | 14 | 22.17 | 3.40 | 4 | 10 |  |  |
| Schizophreniform d/o | 6 | 24.71 | 4.28 | 2 | 4 |  |  |
| Delusional d/o | 1 | 23.17 |  | 1 | 0 |  |  |
| Brief psychotic d/o | 1 | 30.75 |  | 1 | 0 |  |  |
| Psychosis NOS | 3 | 23.41 | 5.33 | 0 | 3 |  |  |
| Major depressive d/o | 20 | 21.50 |  | 12 | 8 |  |  |
| Bipolar disorder | 22 | 24.14 | 4.30 | 8 | 14 |  |  |
| PANSS score |  | Mean Patients | SD Patients | Mean Controls | SD Controls |  |  |
| Total |  | 49.59 | 11.18 | 30.00 | 0.00 |  |  |
| Positive |  | 11.08 | 3.88 | 0.00 | 0.00 |  |  |
| Negative |  | 14.18 | 5.51 | 0.00 | 0.00 |  |  |
| Gen Psychopathology |  | 24.57 | 5.07 | 0.00 | 0.00 |  |  |

### Supplement 8: Non-Affective versus Affective Psychosis

The non-affective psychotic (NAP) disorders are schizophrenia, schizoaffective disorder, schizophreniform disorder, delusional disorder, brief psychotic disorder, and psychosis not otherwise specified. The affective psychotic (AP) disorders are major depressive and Bipolar disorders. Many of the MEM comparisons evaluated in the main text were also examined for differences between NAP and AP.

We examined differences in all MEM metrics among non-affective psychosis (NAP), affective psychosis (AP) patients, and controls using MANCOVA that showed a trend toward significance (Wilk’s λ=0.72, F=1.42, p=0.051). Post hoc between-subjects’ effects for MEM metrics with Bonferroni corrections showed a similar pattern of significant differences between NAP and AP in MEM metrics as psychosis-control comparison, except for the bottom distinct states, which showed a trend toward significance (p=0.08). Another post hoc test to compare NAP, AP, and controls, showed that all MEM metrics with significant differences in the between-subjects’ effects tests were different between controls and NAP but not between controls and AP except top energy trace, which was the highest in NAP, intermediate in AP and the lowest in controls after Games-Howell corrections for multiple testing.

Between NAP and AP patients, no differences in the distributions of distinct states were found in any of the six systems. Between NAP and controls, differences were seen in every system (DMN-top: KS=0.33, p=0.001; DMN-bottom: KS=0.31, p=0.002; DAN-top: KS=0.30, p=0.003; DAN-bottom: KS=0.33, p-0.001; DMN+DAN-top: KS=0.34, p<0.001; DMN+DAN-bottom: KS=0.33, p<0.001). Between AP and controls, significant differences were observed in three systems (DMN-bottom: KS=0.32,p=0.037; DAN-top: KS=0.37, p=0.012; DMN-DAN-top: KS=0.448, p<0.001). NAP always had more distinct states than controls.

Between NAP and AP patients, no differences in the distributions of energy trace were found in any of the six systems. But there were higher energy traces NAP than controls in two systems (DAN-top: KS=0.35, p<0.001; DMN+DAN-top: KS=0.34, p<0.001). Between AP and controls, DMN-DAN-top energy trace was higher in AP (KS=0.32, p<0.038).

Between NAP and AP patients, and between AP and controls, no differences in the distributions of basin transitions states were found in any of the six systems. Between NAP and controls, differences were observed in every system (DMN-top: KS=0.36, p<0.001; DAN-top: KS=0.35,p<0.001; DMN+DAN-top: KS=0.38, p<0.001; DMN+DAN-bottom: KS=0.36, p<0.001) after correcting for multiple tests, where NAP always had more basin transitions than controls.

### Supplement 9: Energy Landscape Results

Basin transition rates for all systems are shown below. For each system, a top graph shows the basin transition matrix, and the bottom graph shows the basin configurations, where the lowest energy basin in the 1^st^ basin in each transition matrix. In each basin transition matrix, each row sums to 100%, and the columns represent time t, while the rows represent time t+1. In each basin configuration plot, the lowest energy is the red basin (colors do not correspond to similar basins between systems; colors just offer a way to identify basins within a system). For each node along the a-axis, black represents “off” while white represents “on.”

It is clear that while the patients have more transitions between basins (as shown in the main text), they do not have different basin transition rates for any of the basins, shown below. In the case of the top switching DMN+DAN, basins 1 and 2 represent states of synchronized DMN+DAN activity, whereas basins 3 and 4 represented asynchrony. Interestingly, for both patients and controls, direct transitions from the DMN+DAN asynchronous basins (3 and 4) is very low (4%), whereas direct transitions from either asynchronous state to a synchronized state are comparatively high (20%).

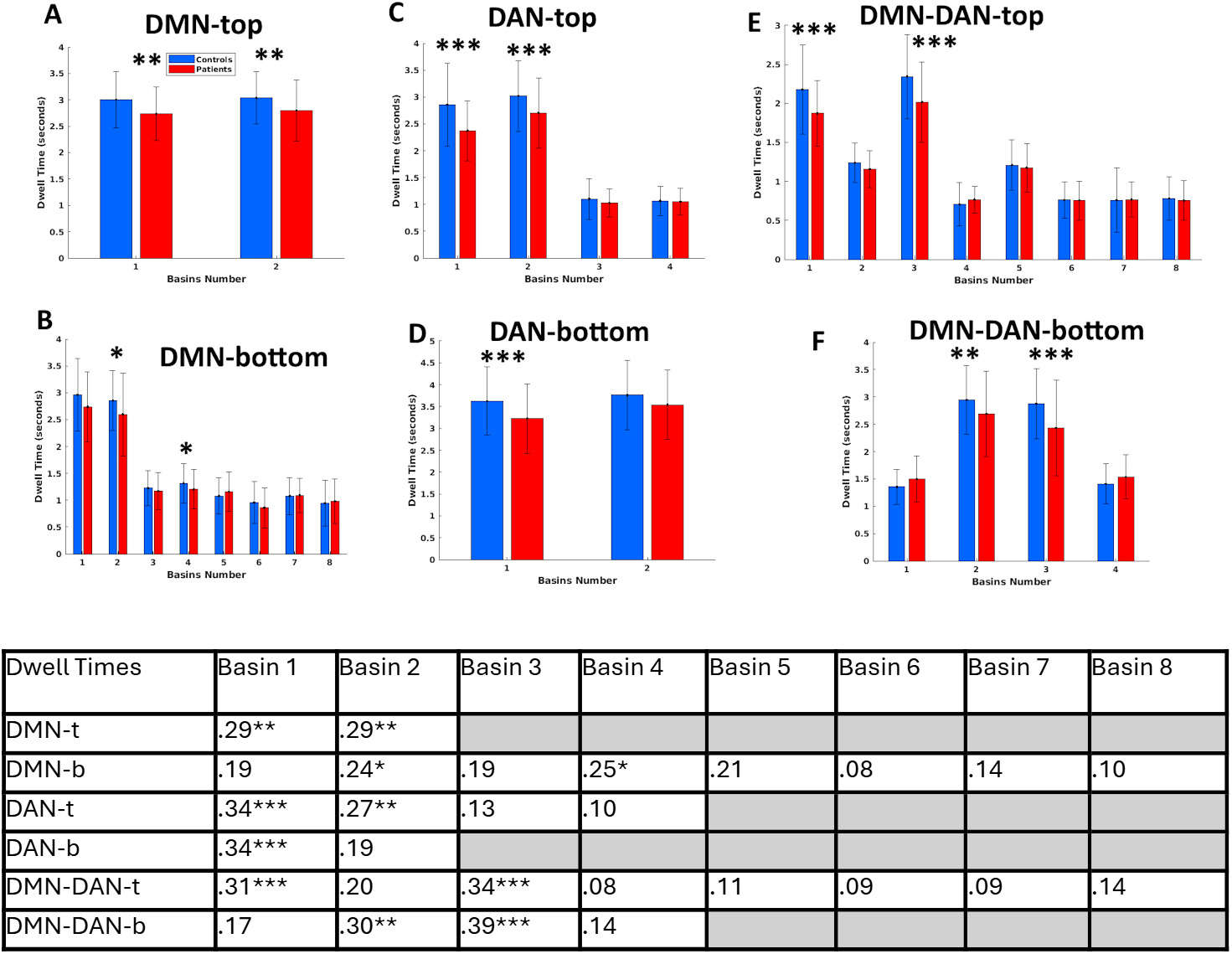
Basin dwell times often differ between groups. **A-F.** Average dwell times for basins associated with all energy minima for controls (blue) and patients (red) for indicated networks. Table shows KS-statistics; asterisks there and above indicate significance of KS-test: *p <0.05, **p <0.01, ***p <0.001. In every case where significance was detected, the patients always had a lower average dwell time for the basin. A variable number of basins was detected per system; where cells are grey (blank), the system did not have that many basins.

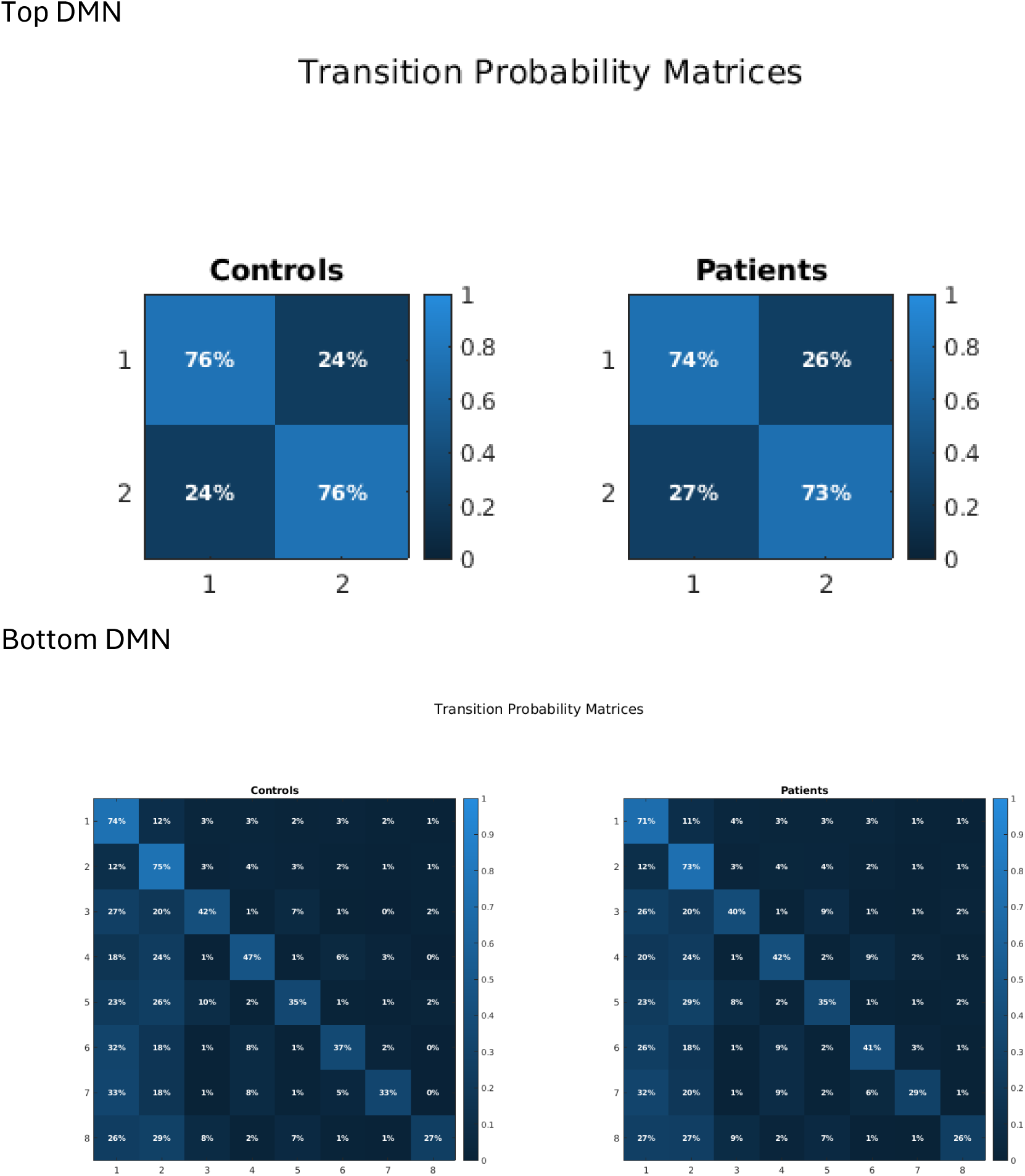
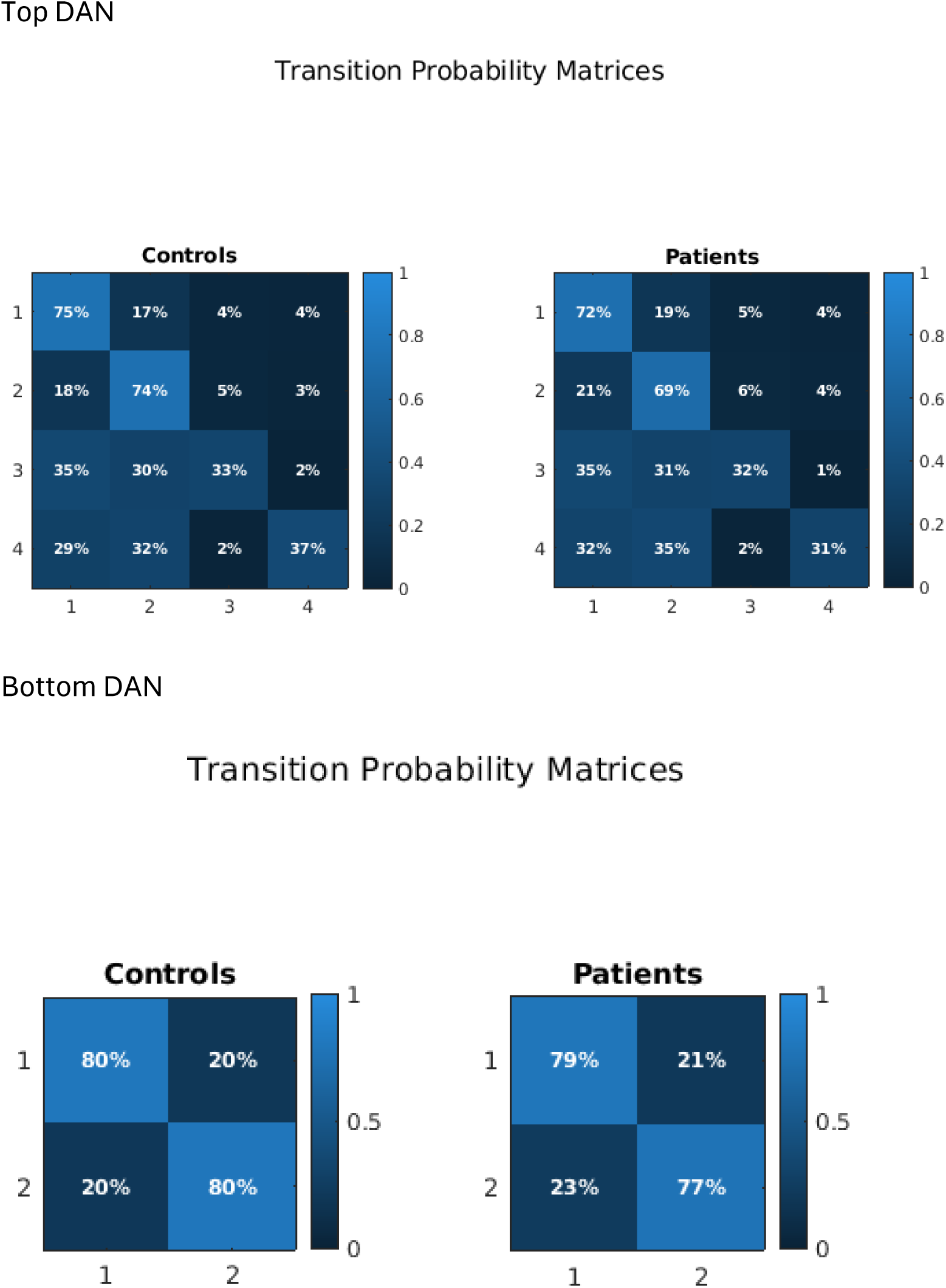
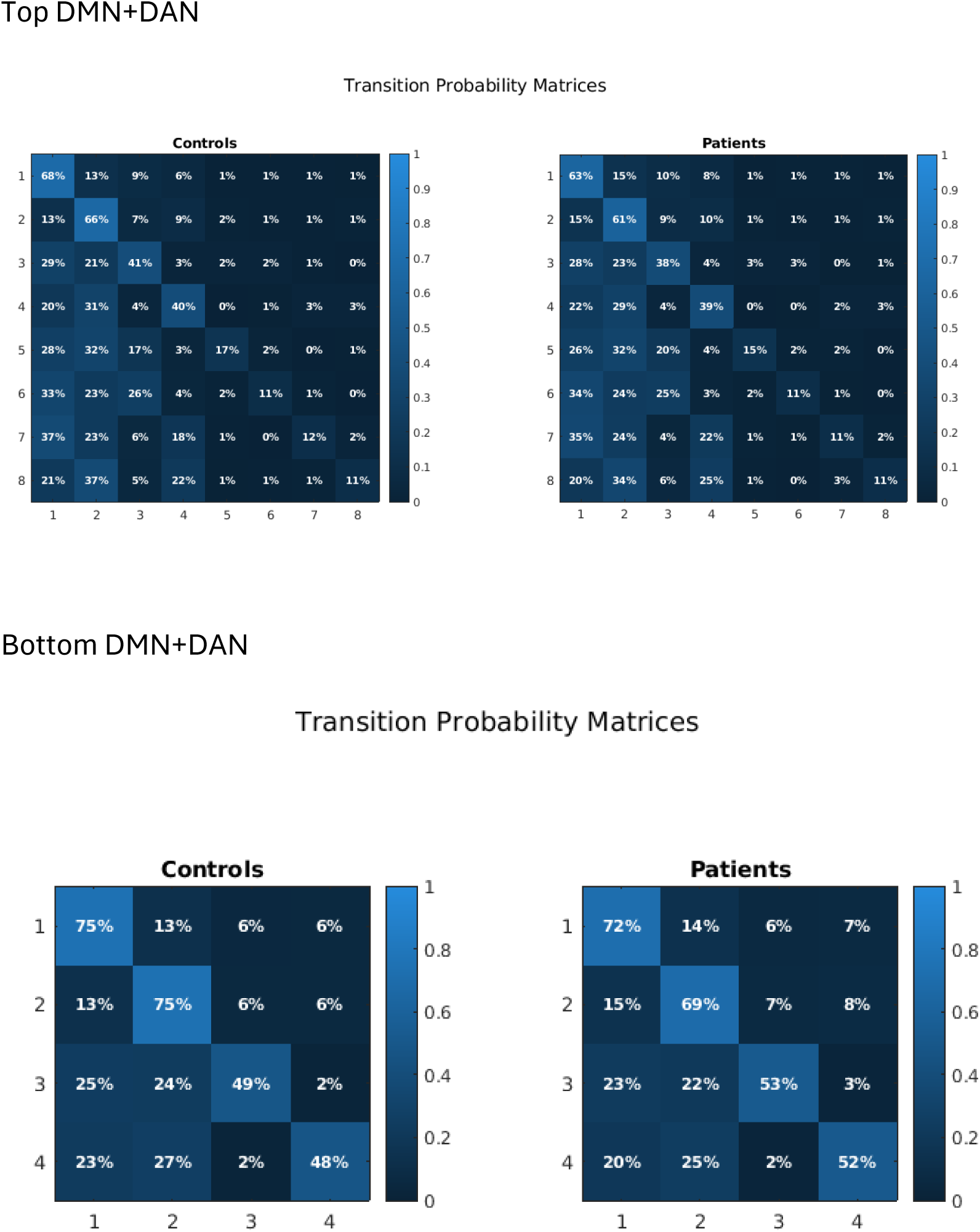

### Supplement 10: Prediction of higher order correlations by MEM

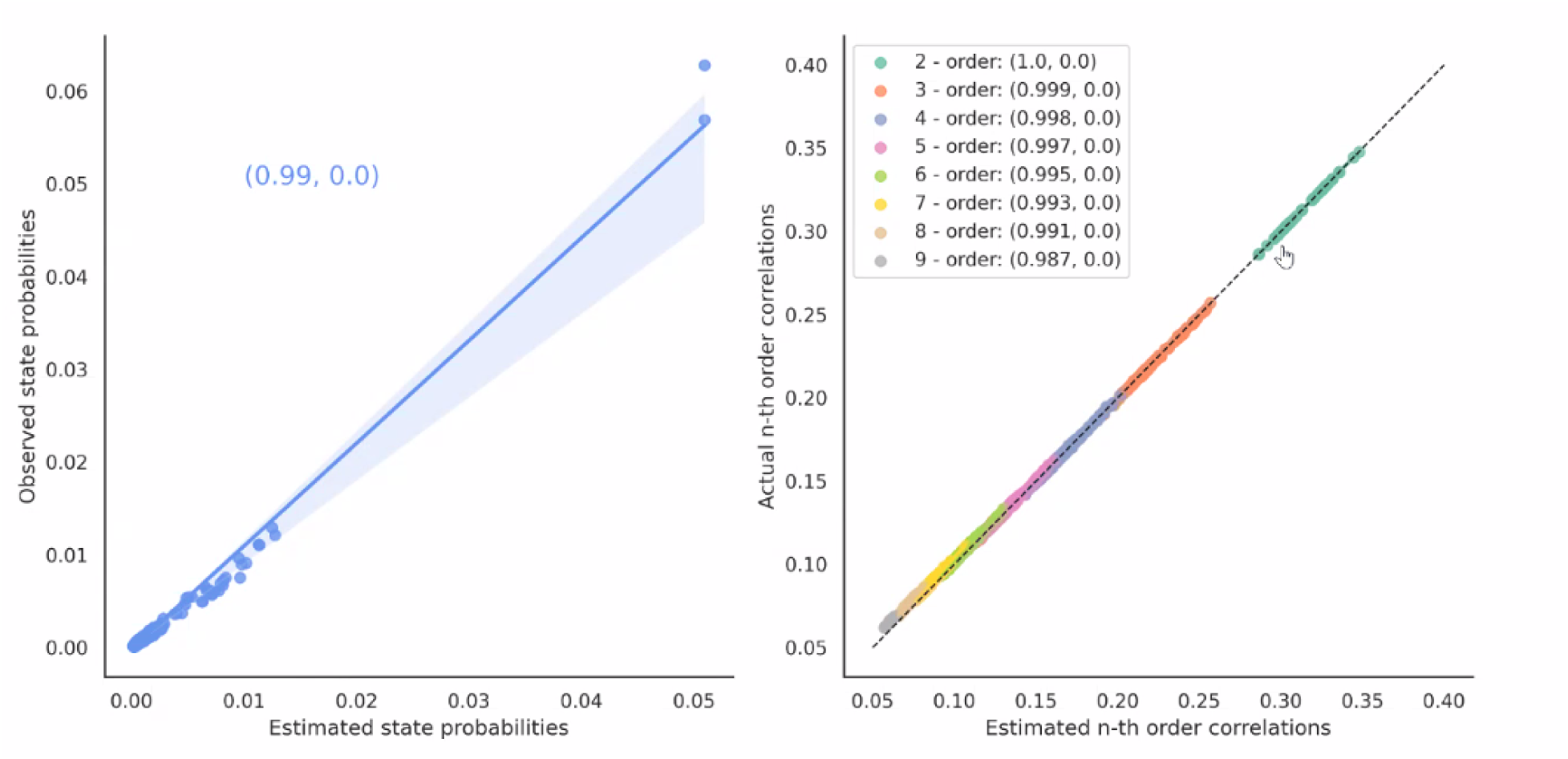

Further, to test whether MEM includes higher order interactions, we examined the correlation between empirical and predicted probabilities of different orders of correlations (from 2^nd^ to the 9^th^ order). For our data, the pairwise MEM accurately reproduced higher-order correlations that were never explicitly fitted indicating that the inferred generative structure provide a sufficiently complete description of the state distribution without requiring explicit higher-order interaction terms. We have now included this analysis for one example network (DMN top) in the revised manuscript

In the figure above, on the left is a scatter plot showing the correlation across 512 unique binary states of the 9-node system. Each state is a scatter point. Two states (the all-on and the all-off states) dominate the probability distribution with a frequency of occurrence of about 0.05. The rest of the states all have probabilities around 0.01 or less. Across all states, the model predicts their occurrence probability with very high accuracy: the correlation between theoretical (model predicted) and empirical (calculated directly from the observations) state probabilities is 0.99.

On the right is a scatter plot showing the correlation of estimated to actual (i.e. empirically observed) “n^th^ order” correlation, where each scatter point shows a combination of nodes for the various combinations according to order (i.e. for 3^rd^ order correlations the temporal similarity of 3 nodes at once). The pairwise MEM model has 1^st^ and 2^nd^ order parameters, yet it still captures higher order parameters.

When n=2 (green scatter points) the range of 2^nd^ order correlations between any two nodes are between 0.26 and 0.35 according to both the model-prediction and empirical data. This shows that for pairwise activity, two nodes are moderately correlated. For higher-order correlations, such as N=3 representing 3-way correlations (orange points), the correlation coefficient is lower, i.e. between 0.25 and 0.2, according to both the model-prediction and empirical data. For even higher order correlations, such as 9^th^ order correlations (grey points) the actual and model-predicted correlations are quite small: barely more than 0.05. Importantly, at every order (pairwise, 3-way, and even up to 9^th^ order), the MEM nearly perfectly explains the observed correlations, and r > 0.98 in every case. Thus, this generative framework can predict the network activation/deactivation globally, primarily using the first-and second-order correlations.

## References

1 O’Neill A, Mechelli A, Bhattacharyya S. Dysconnectivity of Large-Scale Functional Networks in Early Psychosis: A Meta-analysis. Schizophrenia bulletin. 2019;45(3):579–90.

2 Prasad K, Rubin J, Iyengar S, Cape J. Global network disorganization underlying psychosis high risk states. Schizophrenia research. 2023;255:67–68.

3 Lewis M, Santini T, Theis N, Muldoon B, Dash K, Rubin J, et al. Modular architecture and resilience of structural covariance networks in first-episode antipsychotic-naive psychoses. Sci Rep. 2023;13(1):7751.

4 Prasad K, Rubin J, Mitra A, Lewis M, Theis N, Muldoon B, et al. Structural covariance networks in schizophrenia: A systematic review Part II. Schizophrenia research. 2022;239:176–91.

5 Prasad KM, Rubin J, Mitra A, Lewis M, Theis N, Muldoon B, et al. Structural covariance networks in schizophrenia: A systematic review Part I. Schizophrenia research. 2022;240:1–21.

6 Mohanty R, Sethares WA, Nair VA, Prabhakaran V. Rethinking Measures of Functional Connectivity via Feature Extraction. Scientific Reports. 2020;10(1):1298.

7 Reid AT, Headley DB, Mill RD, Sanchez-Romero R, Uddin LQ, Marinazzo D, et al. Advancing functional connectivity research from association to causation. Nature neuroscience. 2019;22(11):1751–60.

8 Hoptman MJ, Zuo XN, Butler PD, Javitt DC, D’Angelo D, Mauro CJ, et al. Amplitude of low-frequency oscillations in schizophrenia: a resting state fMRI study. Schizophrenia research. 2010;117(1):13–20.

9 Amaro E, Jr., Barker GJ. Study design in fMRI: basic principles. Brain Cogn. 2006;60(3):220–32.

10 Keshavan MS, Eack SM, Prasad KM, Haller CS, Cho RY. Longitudinal functional brain imaging study in early course schizophrenia before and after cognitive enhancement therapy. NeuroImage. 2017;151:55–64.

11 Stolz E, Pancholi KM, Goradia DD, Paul S, Keshavan MS, Nimgaonkar VL, et al. Brain activation patterns during visual episodic memory processing among first-degree relatives of schizophrenia subjects. NeuroImage. 2012;63(3):1154–61.

12 Glahn DC, Ragland JD, Abramoff A, Barrett J, Laird AR, Bearden CE, et al. Beyond hypofrontality: a quantitative meta-analysis of functional neuroimaging studies of working memory in schizophrenia. Human brain mapping. 2005;25(1):60–9.

13 Ragland JD, Laird AR, Ranganath C, Blumenfeld RS, Gonzales SM, Glahn DC. Prefrontal activation deficits during episodic memory in schizophrenia. The American journal of psychiatry. 2009;166(8):863–74.

14 Cai M, Wang R, Liu M, Du X, Xue K, Ji Y, et al. Disrupted local functional connectivity in schizophrenia: An updated and extended meta-analysis. Schizophrenia. 2022;8(1):93.

15 Prasad KM, Muldoon B, Theis N, Iyengar S, Keshavan MS. Multipronged investigation of morphometry and connectivity of hippocampal network in relation to risk for psychosis using ultrahigh field MRI. Schizophrenia research. 2023;256:88–97.

16 Sanfratello L, Houck JM, Calhoun VD. Dynamic Functional Network Connectivity in Schizophrenia with Magnetoencephalography and Functional Magnetic Resonance Imaging: Do Different Timescales Tell a Different Story? Brain Connect. 2019;9(3):251–62.

17 Damaraju E, Allen EA, Belger A, Ford JM, McEwen S, Mathalon DH, et al. Dynamic functional connectivity analysis reveals transient states of dysconnectivity in schizophrenia. NeuroImage Clinical. 2014;5:298–308.

18 Wang X, Chang Z, Wang R. Opposite effects of positive and negative symptoms on resting-state brain networks in schizophrenia. Commun Biol. 2023;6(1):279.

19 Matkovič A, Anticevic A, Murray JD, Repovš G. Static and dynamic fMRI-derived functional connectomes represent largely similar information. Network Neuroscience. 2023;7(4):1266–301.

20 Masuda N, Islam S, Thu Aung S, Watanabe T. Energy landscape analysis based on the Ising model: Tutorial review. PLOS Complex Systems. 2025;2(5):e0000039.

21 Prasad K, Bowei O, Theis N. From microstates to macroscales: A Critical Review of Maximum Entropy Modeling and Energy Landscape Analysis in functional MRI. Res Sq. 2025.

22 Prasad K, Bowei O, Theis N. From microstates to macroscales: A critical review of maximum entropy modeling and energy landscape analysis in functional MRI. Neurosci Biobehav Rev. 2026;188:106834.

23 Ishida T, Yamada S, Yasuda K, Uenishi S, Tamaki A, Tabata M, et al. Aberrant brain dynamics of large-scale functional networks across schizophrenia and mood disorder. NeuroImage: Clinical. 2024;41:103574.

24 Jacobs GR, Coleman MJ, Lewandowski KE, Pasternak O, Cetin-Karayumak S, Mesholam-Gately RI, et al. An Introduction to the Human Connectome Project for Early Psychosis. Schizophrenia bulletin. 2025;51(3):658–71.

25 Dafflon J, F. Da Costa P, Váša F, Monti RP, Bzdok D, Hellyer PJ, et al. A guided multiverse study of neuroimaging analyses. Nature Communications. 2022;13(1):3758.

26 Steegen S, Tuerlinckx F, Gelman A, Vanpaemel W. Increasing Transparency Through a Multiverse Analysis. Perspect Psychol Sci. 2016;11(5):702–12.

27 Gohel SR, Biswal BB. Functional integration between brain regions at rest occurs in multiple-frequency bands. Brain Connect. 2015;5(1):23–34.

28 Bluhm RL, Miller J, Lanius RA, Osuch EA, Boksman K, Neufeld RW, et al. Spontaneous low-frequency fluctuations in the BOLD signal in schizophrenic patients: anomalies in the default network. Schizophrenia bulletin. 2007;33(4):1004–12.

29 Hirano Y, Oribe N, Kanba S, Onitsuka T, Nestor PG, Spencer KM. Spontaneous Gamma Activity in Schizophrenia. JAMA psychiatry. 2015;72(8):813–21.

30 Garrity AG, Pearlson GD, McKiernan K, Lloyd D, Kiehl KA, Calhoun VD. Aberrant “default mode” functional connectivity in schizophrenia. The American journal of psychiatry. 2007;164(3):450–7.

31 Menon V. 20 years of the default mode network: A review and synthesis. Neuron. 2023;111(16):2469–87.

32 Voineskos AN, Hawco C, Neufeld NH, Turner JA, Ameis SH, Anticevic A, et al. Functional magnetic resonance imaging in schizophrenia: current evidence, methodological advances, limitations and future directions. World Psychiatry. 2024;23(1):26–51.

33 Finn ES. Is it time to put rest to rest? Trends in cognitive sciences. 2021;25(12):1021–32.

34 Esposito R, Cieri F, Chiacchiaretta P, Cera N, Lauriola M, Di Giannantonio M, et al. Modifications in resting state functional anticorrelation between default mode network and dorsal attention network: comparison among young adults, healthy elders and mild cognitive impairment patients. Brain imaging and behavior. 2018;12(1):127–41.

35 Owens MM, Yuan D, Hahn S, Albaugh M, Allgaier N, Chaarani B, et al. Investigation of Psychiatric and Neuropsychological Correlates of Default Mode Network and Dorsal Attention Network Anticorrelation in Children. Cereb Cortex. 2020;30(12):6083–96.

36 Raichle ME. The restless brain: how intrinsic activity organizes brain function. Philosophical transactions of the Royal Society of London Series B, Biological sciences. 2015;370(1668).

37 Martin AK, Mowry B, Reutens D, Robinson GA. Executive functioning in schizophrenia: Unique and shared variance with measures of fluid intelligence. Brain Cogn. 2015;99:57–67.

38 Theis N, Bahuguna J, Rubin JE, Banerjee SS, Muldoon B, Prasad KM. Energy of Functional Brain States Correlates With Cognition in Adolescent-Onset Schizophrenia and Healthy Persons. Human brain mapping. 2025;46(1):e70129.

39 Glasser MF, Coalson TS, Robinson EC, Hacker CD, Harwell J, Yacoub E, et al. A multi-modal parcellation of human cerebral cortex. Nature. 2016;536(7615):171–78.

40 Glasser MF, Sotiropoulos SN, Wilson JA, Coalson TS, Fischl B, Andersson JL, et al. The minimal preprocessing pipelines for the Human Connectome Project. NeuroImage. 2013;80:105–24.

41 Buitrago P, Nystrom N. 205–19 (High Performance Computing, 2021).

42 Fischl B. FreeSurfer. NeuroImage. 2012;62(2):774–81.

43 Jenkinson M, Beckmann CF, Behrens TE, Woolrich MW, Smith SM. Fsl. NeuroImage. 2012;62(2):782–90.

44 Holmes A, Levi PT, Chen YC, Chopra S, Aquino KM, Pang JC, et al. Disruptions of Hierarchical Cortical Organization in Early Psychosis and Schizophrenia. Biol Psychiatry Cogn Neurosci Neuroimaging. 2023;8(12):1240–50.

45 Lewis M, Theis N, Girish N, Prasad K. Determining the atlas correspondence of Desikan-Killiany-Tourville and Glasser MMP1 atlases across magnetic field strengths. Journal of neuroscience methods. 2025;418:110445.

46 Sandhu Z, Tanglay O, Young IM, Briggs RG, Bai MY, Larsen ML, et al. Parcellation-based anatomic modeling of the default mode network. Brain Behav. 2021;11(2):e01976.

47 Allan PG, Briggs RG, Conner AK, O’Neal CM, Bonney PA, Maxwell BD, et al. Parcellation-based tractographic modeling of the dorsal attention network. Brain Behav. 2019;9(10):e01365.

48 Zuo XN, Di Martino A, Kelly C, Shehzad ZE, Gee DG, Klein DF, et al. The oscillating brain: complex and reliable. NeuroImage. 2010;49(2):1432–45.

49 Epp SM, Castrillón G, Yuan B, Andrews-Hanna J, Preibisch C, Riedl V. BOLD signal changes can oppose oxygen metabolism across the human cortex. Nature neuroscience. 2025.

50 Kennerley AJ, Harris S, Bruyns-Haylett M, Boorman L, Zheng Y, Jones M, et al. Early and late stimulus-evoked cortical hemodynamic responses provide insight into the neurogenic nature of neurovascular coupling. Journal of cerebral blood flow and metabolism : official journal of the International Society of Cerebral Blood Flow and Metabolism. 2012;32(3):468–80.

51 Drew PJ. Vascular and neural basis of the BOLD signal. Curr Opin Neurobiol. 2019;58:61–69.

52 Penttonen M, Buzsáki G. Natural logarithmic relationship between brain oscillators. Thalamus & Related Systems. 2003;2:145–52.

53 Logothetis NK. The Underpinnings of the BOLD Functional Magnetic Resonance Imaging Signal. The Journal of Neuroscience. 2003;23(10):3963.

54 Birn RM, Diamond JB, Smith MA, Bandettini PA. Separating respiratory-variation-related fluctuations from neuronal-activity-related fluctuations in fMRI. NeuroImage. 2006;31(4):1536–48.

55 Moon HS, Jiang H, Vo TT, Jung WB, Vazquez AL, Kim SG. Contribution of Excitatory and Inhibitory Neuronal Activity to BOLD fMRI. Cereb Cortex. 2021;31(9):4053–67.

56 Rubinov M, Kötter R, Hagmann P, Sporns O. Brain connectivity toolbox: a collection of complex network measurements and brain connectivity datasets. NeuroImage. 2009;47:S169.

57 Liu DC, Nocedal J. On the Limited Memory Bfgs Method for Large-Scale Optimization. Mathematical Programming. 1989;45(3):503–28.

58 Cho JW, Korchmaros A, Vogelstein JT, Milham MP, Xu T. Impact of concatenating fMRI data on reliability for functional connectomics. NeuroImage. 2021;226:117549.

59 Watanabe T, Hirose S, Wada H, Imai Y, Machida T, Shirouzu I, et al. A pairwise maximum entropy model accurately describes resting-state human brain networks. Nat Commun. 2013;4:1370.

60 Fu Z, Iraji A, Sui J, Calhoun VD. Whole-Brain Functional Network Connectivity Abnormalities in Affective and Non-Affective Early Phase Psychosis. Frontiers in Neuroscience. 2021;Volume 15 -2021.

61 Wang HR, Liu ZQ, Nakua H, Hegarty CE, Thies MB, Patel PK, et al. Decoding Early Psychoses: Unraveling Stable Microstructural Features Associated With Psychopathology Across Independent Cohorts. Biological psychiatry. 2025;97(2):167–77.

62 O’Neill AG, Pax M, Parent JH, Sepulcre J, Camprodon JA, Noble S, et al. Predicting cognitive functioning in early psychosis: factors supporting and limiting generalizability of connectome-based models. NPP—Digital Psychiatry and Neuroscience. 2025;3(1):11.

63 Meda SA, Wang Z, Ivleva EI, Poudyal G, Keshavan MS, Tamminga CA, et al. Frequency-Specific Neural Signatures of Spontaneous Low-Frequency Resting State Fluctuations in Psychosis: Evidence From Bipolar-Schizophrenia Network on Intermediate Phenotypes (B-SNIP) Consortium. Schizophrenia bulletin. 2015;41(6):1336–48.

64 Gohel S, Gallego JA, Robinson DG, DeRosse P, Biswal B, Szeszko PR. Frequency specific resting state functional abnormalities in psychosis. Human brain mapping. 2018;39(11):4509–18.

65 Logothetis NK. The ins and outs of fMRI signals. Nature neuroscience. 2007;10(10):1230–2.

66 Murray JD, Wang XJ. Cortical circuit models in psychiatry: Linking disrupted ecitation-inhibition balance to cognitive deficits associated with schizophrenia. In: Anticevic A, Murray JD, editors. Computational Psychiatry: Mathematical Modeling of Mental Illness. Cambridge, MA: Academic Press; 2018. p. 3–29.

67 Deco G, Jirsa VK, McIntosh AR. Emerging concepts for the dynamical organization of resting-state activity in the brain. Nature reviews Neuroscience. 2011;12(1):43–56.

68 Mueller S, Wang D, Fox MD, Yeo BT, Sepulcre J, Sabuncu MR, et al. Individual variability in functional connectivity architecture of the human brain. Neuron. 2013;77(3):586–95.

69 Simoncelli EP, Olshausen BA. Natural image statistics and neural representation. Annu Rev Neurosci. 2001;24:1193–216.

70 Fuhs MC, Touretzky DS. A spin glass model of path integration in rat medial entorhinal cortex. The Journal of neuroscience : the official journal of the Society for Neuroscience. 2006;26(16):4266–76.

71 Tang A, Jackson D, Hobbs J, Chen W, Smith JL, Patel H, et al. A maximum entropy model applied to spatial and temporal correlations from cortical networks in vitro. The Journal of neuroscience : the official journal of the Society for Neuroscience. 2008;28(2):505–18.

72 Das TK, Abeyasinghe PM, Crone JS, Sosnowski A, Laureys S, Owen AM, et al. Highlighting the structure-function relationship of the brain with the Ising model and graph theory. Biomed Res Int. 2014;2014:237898.

73 Ezaki T, Watanabe T, Ohzeki M, Masuda N. Energy landscape analysis of neuroimaging data. Philos Trans A Math Phys Eng Sci. 2017;375(2096):20160287.

74 Gu S, Cieslak M, Baird B, Muldoon SF, Grafton ST, Pasqualetti F, et al. The Energy Landscape of Neurophysiological Activity Implicit in Brain Network Structure. Scientific Reports. 2018;8(1):2507.

75 Kang J, Jeong SO, Pae C, Park HJ. Bayesian estimation of maximum entropy model for individualized energy landscape analysis of brain state dynamics. Human brain mapping. 2021;42(11):3411–28.

76 Savin C, Tkačik G. Maximum entropy models as a tool for building precise neural controls. Curr Opin Neurobiol. 2017;46:120–26.

77 Amit DJ, Gutfreund H, Sompolinsky H. Spin-glass models of neural networks. Phys Rev A Gen Phys. 1985;32(2):1007–18.

78 Yeh F-C, Tang A, Hobbs JP, Hottowy P, Dabrowski W, Sher A, et al. Maximum Entropy Approaches to Living Neural Networks. Entropy. 2010;12(1):89–106.

79 Bowei O, Theis N, Rubin J, Prasad KM. Dynamic network reconfigurations during task engagement following resting state in adolescent onset schizophrenia. bioRxiv. 2026:2026.06.05.730513.

80 Kindler J, Ishida T, Michel C, Klaassen AL, Stuble M, Zimmermann N, et al. Aberrant brain dynamics in individuals with clinical high risk of psychosis. Schizophr Bull Open. 2024;5(1).

81 Das A, de Los Angeles C, Menon V. Electrophysiological foundations of the human default-mode network revealed by intracranial-EEG recordings during resting-state and cognition. NeuroImage. 2022;250:118927.

82 Corbetta M, Shulman GL. Control of goal-directed and stimulus-driven attention in the brain. Nature Reviews Neuroscience. 2002;3(3):201–15.

83 Fox MD, Corbetta M, Snyder AZ, Vincent JL, Raichle ME. Spontaneous neuronal activity distinguishes human dorsal and ventral attention systems. Proceedings of the National Academy of Sciences. 2006;103(26):10046–51.

84 Pagani M, Gutierrez-Barragan D, de Guzman AE, Xu T, Gozzi A. Mapping and comparing fMRI connectivity networks across species. Commun Biol. 2023;6(1):1238.

85 Gutierrez-Barragan D, Ramirez JSB, Panzeri S, Xu T, Gozzi A. Evolutionarily conserved fMRI network dynamics in the mouse, macaque, and human brain. Nature Communications. 2024;15(1):8518.

86 Spreng RN, Stevens WD, Chamberlain JP, Gilmore AW, Schacter DL. Default network activity, coupled with the frontoparietal control network, supports goal-directed cognition. NeuroImage. 2010;53(1):303–17.

87 Whitfield-Gabrieli S, Ford JM. Default mode network activity and connectivity in psychopathology. Annu Rev Clin Psychol. 2012;8:49–76.

88 Lavigne KM, Menon M, Woodward TS. Functional Brain Networks Underlying Evidence Integration and Delusions in Schizophrenia. Schizophrenia bulletin. 2020;46(1):175–83.

89 Hur JW, Kim T, Cho KIK, Kwon JS. Attenuated Resting-State Functional Anticorrelation between Attention and Executive Control Networks in Schizotypal Personality Disorder. J Clin Med. 2021;10(2).

90 Green MF, Kern RS, Braff DL, Mintz J. Neurocognitive deficits and functional outcome in schizophrenia: are we measuring the “right stuff”? Schizophrenia bulletin. 2000;26(1):119–36.

91 Elvevag B, Goldberg TE. Cognitive impairment in schizophrenia is the core of the disorder. Critical reviews in neurobiology. 2000;14(1):1–21.

92 Eack SM, Greenwald DP, Hogarty SS, Keshavan MS. One-year durability of the effects of cognitive enhancement therapy on functional outcome in early schizophrenia. Schizophrenia research. 2010;120(1-3):210–6.

93 Eack SM, Hogarty GE, Cho RY, Prasad KM, Greenwald DP, Hogarty SS, et al. Neuroprotective effects of cognitive enhancement therapy against gray matter loss in early schizophrenia: results from a 2-year randomized controlled trial. Archives of general psychiatry. 2010;67(7):674–82.

## REFERENCES

1. Liu DC, Nocedal J (1989): On the Limited Memory Bfgs Method for Large-Scale Optimization. Mathematical Programming. 45:503–528.

